# Hog1/p38 regulates the integrated stress response (ISR) and global transcriptomic changes in *C. neoformans* during oxidative stress

**DOI:** 10.1101/2025.05.19.654802

**Authors:** David Goich, Julia R. Furfaro, John C. Panepinto

**Author notes:** Corresponding author: John C. Panepinto.

## Abstract

*Cryptococcus neoformans* is an opportunistic fungal pathogen that is responsible for an estimated 112,000 deaths per year. We studied the role of the Hog1/p38 and Gcn2 signal transduction pathways during adaptation of *C. neoformans* to oxidative stress, a physiologically relevant stressor during infection. Using a combination of molecular assays and RNA-sequencing analysis in Hog1-, Gcn2-and double-deleted mutants, we identify shared targets of these pathways, and investigate a negative feedback loop that regulates induction of the integrated stress response (ISR). We found that simultaneous loss of Hog1 and Gcn2 profoundly impacts the ability of *C. neoformans* to tolerate oxidative stress. We also show that Hog1 regulates induction of the ISR, and that the Hog1 and Gcn2 pathways converge on repression of abundant, pro-growth mRNA, a key step in translatome reprogramming. Our results establish Hog1 and Gcn2 as key regulators of the response to oxidative stress, and demonstrate that the interplay between these pathways is critical for ROS tolerance in *C. neoformans*. More broadly, our results demonstrate the extent to which signaling via Hog1/p38 and Gcn2 are linked, and point to extensive recalibration of the cellular signaling apparatus when one of these pathways is disrupted.

**Author Summary:** *Cryptococcus neoformans* is a fungus that causes deadly infections, resulting in approximately 112,000 deaths per year. In this study, we examined two genes that are important for the ability of *C. neoformans* to tolerate stressors it would encounter in the human body: Hog1 and Gcn2. We found that Hog1 and Gcn2 perform overlapping functions during the fungal response to stress. These findings are important because they highlight the potential benefits of targeting these pathways in conjunction to treat infection with *C. neoformans*. These findings could be used to develop new drug or treatment strategies to reduce the disease burden associated with this infection.

## Introduction

*Cryptococcus neoformans* is a ubiquitous environmental fungus and opportunistic pathogen of people. Cryptococcosis, infection with *Cryptococcus*, is responsible for an estimated 112,000 deaths per year, primarily in people who are immunocompromised (1). Cryptococcosis usually begins with primary infection of the lungs, which can then disseminate to the central nervous system (CNS) if there are defects in pulmonary containment (2). Adaptation to the host environment is an important prerequisite for virulence of systemic fungal pathogens such as *C. neoformans*. Thus, understanding the mechanisms which mitigate host-derived stress is key to our understanding of how fungi achieve virulence.

A major strategy employed by the host to restrict growth of invading microbes involves production of reactive oxygen and nitrogen species (ROS and RNS) by the host immune system, reviewed in (3).

These strategies are associated with the oxidative burst, which occurs inside the phagosomes of innate immune cells such as macrophages and neutrophils to kill invading microbes. Many canonical pathways of reducing ROS are present in *C. neoformans*, such as glutathione (GSH) and thioredoxin (Trx) systems, as well as catalases and superoxide dismutases (SODs) (4–7). In addition to directly neutralizing ROS, fungi must also upregulate repair and degradation pathways to ensure that damaged biomolecules do not disrupt cellular homeostasis. Given the ability of ROS to produce broad-spectrum of damage to macromolecules such as lipids (8), proteins (9), and nucleic acids (10), the response of pathogenic fungi such as *C. neoformans* to oxidative stress is complex and multifaceted..

Successful adaptation to stress necessitates a coordinated response which requires reallocation of translational capacity toward stress-related mRNAs (11). This process, which we call **translatome reprogramming**, is a conserved component of stress responses and has a few major features: (i) Production of stress-mitigating proteins is increased. (ii) The abundance of homeostatic, pro-growth mRNA is reduced. This category of transcripts often encodes components of the ribosome (12–14), which are some of the most abundant transcripts in the unstressed cell. (iii) Translation is repressed (15), which frees pro-growth mRNA for decay and frees ribosomes to associate with stress-related mRNA.

Translatome reprogramming is regulated by signal transduction pathways, such as the Gcn2 pathway. Gcn2 is a kinase activated by stress which targets translation initiation factor 2α (eIF2α) for phosphorylation (16). Phosphorylated eIF2α binds and inhibits initiation factor 2B (eIF2B), preventing its activity as a guanine nucleotide exchange factor (GEF) in ternary complex formation (17). This depletion of ternary complex results in global repression of cap-dependent translation initiation (18). Gcn2 is activated by uncharged tRNAs and ribosome collisions, which are stress-derived signals (19). In the case of ROS, oxidative damage to mRNAs, ribosomes, or tRNAs could result in ribosome stalling and collisions (20, 21), or the inability of tRNAs to be charged.

While Gcn2 activity represses translation at a global scale, it also promotes selective translation of translationally regulated mRNAs, such as the *GCN4* mRNA (22). While the mechanism of *GCN4* translation is not fully understood, depletion of ternary complex promotes bypass of inhibitory upstream open reading frames (uORFs) in the *GCN4* mRNA, resulting in translation of the main coding sequence of this transcript (23). Gcn4 is the transcription factor (TF) responsible for the integrated stress response (ISR), which upregulates transcripts related to amino acid biosynthesis and NADPH-dependent oxidoreductases, which can shift metabolic flux toward a reducing environment (13, 24). We’ve previously demonstrated that the Gcn2 pathway regulates several aspects of translatome reprogramming in *C. neoformans* during oxidative stress, and that strains lacking Gcn2 have reduced oxidative stress tolerance (13, 15).

Another important pathway in the response to oxidative stress in *C. neoformans* is the Hog1/p38 pathway (25). Hog1 is a p38 MAP kinase (MAPK) that is activated in response to a broad spectrum of stressors, and is known to regulate the transcriptional, post-transcriptional, and translational response to stress in other fungi (26). While comparatively less is known about how Hog1 is activated by oxidative stress and its direct targets in *C. neoformans*, it is nonetheless important for ROS tolerance, since Hog1 deletion mutants have reduced survival upon exposure to oxidants such as hydrogen peroxide (25).

Recently, we demonstrated that simultaneous engagement of the Hog1 and Gcn2 pathways during thermal stress was associated with crosstalk between these pathways (27). Notably, loss of Hog1 was associated with increased output via the Gcn2 pathway, and this compensatory signaling appeared to promote thermoadaptation. In the present study, we extend this area of research to oxidative stress, and find that while these pathways also undergo crosstalk in the oxidative stress response, the mechanism appears to be distinct.

By phenotypic analysis, we find that simultaneous disruption of Hog1 and Gcn2 results in profound oxidative stress sensitivity. Using polysome profiling and western blotting, we find that Hog1 regulates ribosome reallocation and Gcn4 levels in response to oxidative stress without altering the amount of P-eIF2α. To better understand the contributions of Hog1 and Gcn2 to oxidative stress response, we performed mRNA-sequencing of Hog1-, Gcn2-, and double-deletion strains. We found that while Hog1 deletion causes a transcriptome-wide defect in the oxidative stress response, the ISR is robustly upregulated in this strain. We also demonstrate that both pathways regulate the repression of pro-growth transcripts that encode ribosome-associated factors, and that disruption of both Hog1 and Gcn2 prevents downregulation of these transcripts.

Overall, our findings indicate that Hog1 and Gcn2 play a major role in the sensitivity of *C. neoformans* to oxidative stress, and profoundly impact translation and the translating pool of mRNAs in the response to ROS. Our results suggest that compensatory induction of the Gcn2-dependent ISR in *hog1*Δ promotes its survival during oxidative stress despite large-scale defects in its response to this stressor. This work also has important implications in the study of signal transduction, demonstrating that disruption of core signal transduction pathways can globally recalibrate homeostasis and stress responses, pointing to the difficulty of studying these pathways in isolation.

## Results

### Simultaneous loss of Hog1 and Gcn2 exacerbates defects in oxidative stress tolerance

Previously, we demonstrated that the Hog1 and Gcn2 pathways converge on the process of translatome reprogramming during thermal stress, and that simultaneous disruption of these pathways results in exacerbated thermosensitivity (27). Thus, we initially sought to determine whether Hog1 and Gcn2 played a similar role in regulating sensitivity to oxidative stress. To test this, we performed spot plate analysis of wild-type (WT), *hog1*Δ, *gcn2*Δ, and *hog1*Δ*gcn2*Δ strains grown in the presence or absence of hydrogen peroxide (**Fig. 1**). We found that while *hog1*Δ and *gcn2*Δ exhibited some sensitivity to oxidative stress, *hog1*Δ*gcn2*Δ double mutants were profoundly sensitive to growth in the presence of H_2_O_2_. Given the greatly exacerbated sensitivity of the double mutant, these results provide evidence of a negative genetic interaction between Hog1 and Gcn2 in the context of ROS.

**Figure 1:**
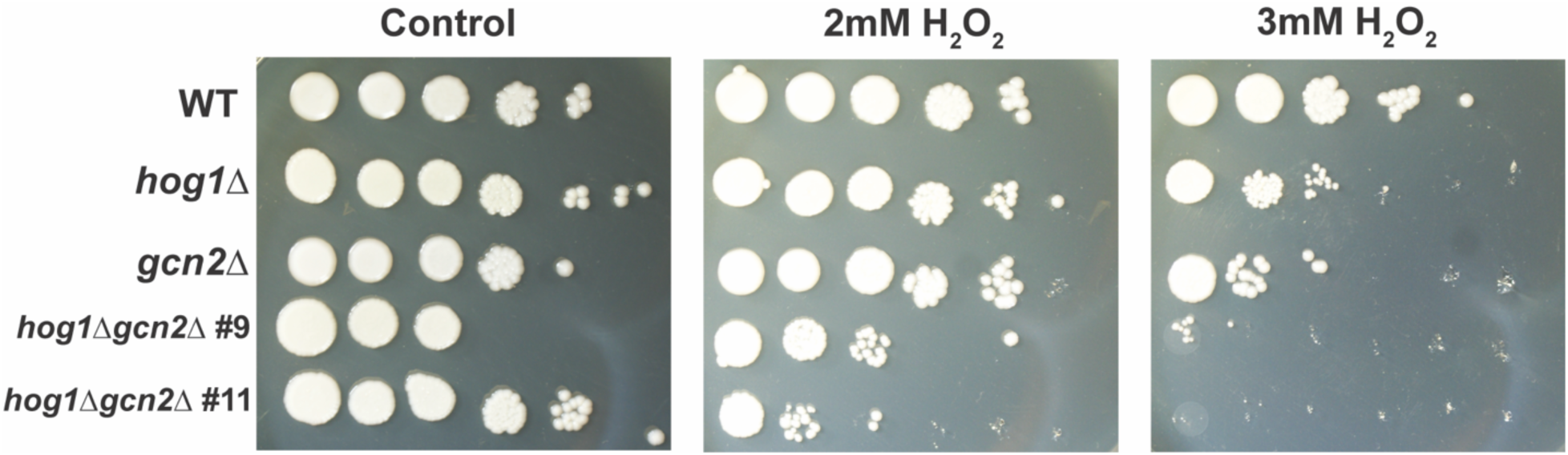
Loss of Hog1 and Gcn2 exacerbates sensitivity to ROS. WT, *hog1*Δ, *gcn2*Δ, and two clones of *hog1*Δ*gcn2*Δ were grown overnight and serially diluted for spot plate analysis. Cells were subjected to oxidative stress by incubation on plates containing the indicated concentration of hydrogen peroxide. A representative biological replicate is shown (*n = 3*).

### Loss of Hog1 is associated with decreased translation repression in response to ROS

We next began to investigate the nature of the genetic interaction between Hog1 and Gcn2 during the response to oxidative stress. We previously found that Hog1 deletion was associated with increased translation repression in response to thermal stress via the Gcn2 pathway (27). Thus, we set out to recapitulate this finding in the context of oxidative stress. To test this, we performed polysome profiling of actively dividing WT and *hog1*Δ cells in response to oxidative stress, which was simulated using 2mM H_2_O_2_ (**Fig. 2A**). In the WT response, we observed robust repression of translation, as evidenced by collapse of polysome peaks and robust accumulation of free 60S ribosomal subunits (**Fig. 2A**, left). In *hog1*Δ, translation repression also occurred but appeared reduced in its extent, with persistence of polysomes after thirty minutes of oxidative stress (**Fig. 2A**, right). Given our previous findings in the context of thermal stress (27), this result was surprising, and indicated that the effect of Hog1 deletion on translational response is stress-specific. In the context of oxidative stress, our data suggest that Hog1 deletion results in a dampened translational response.

**Figure 2:**
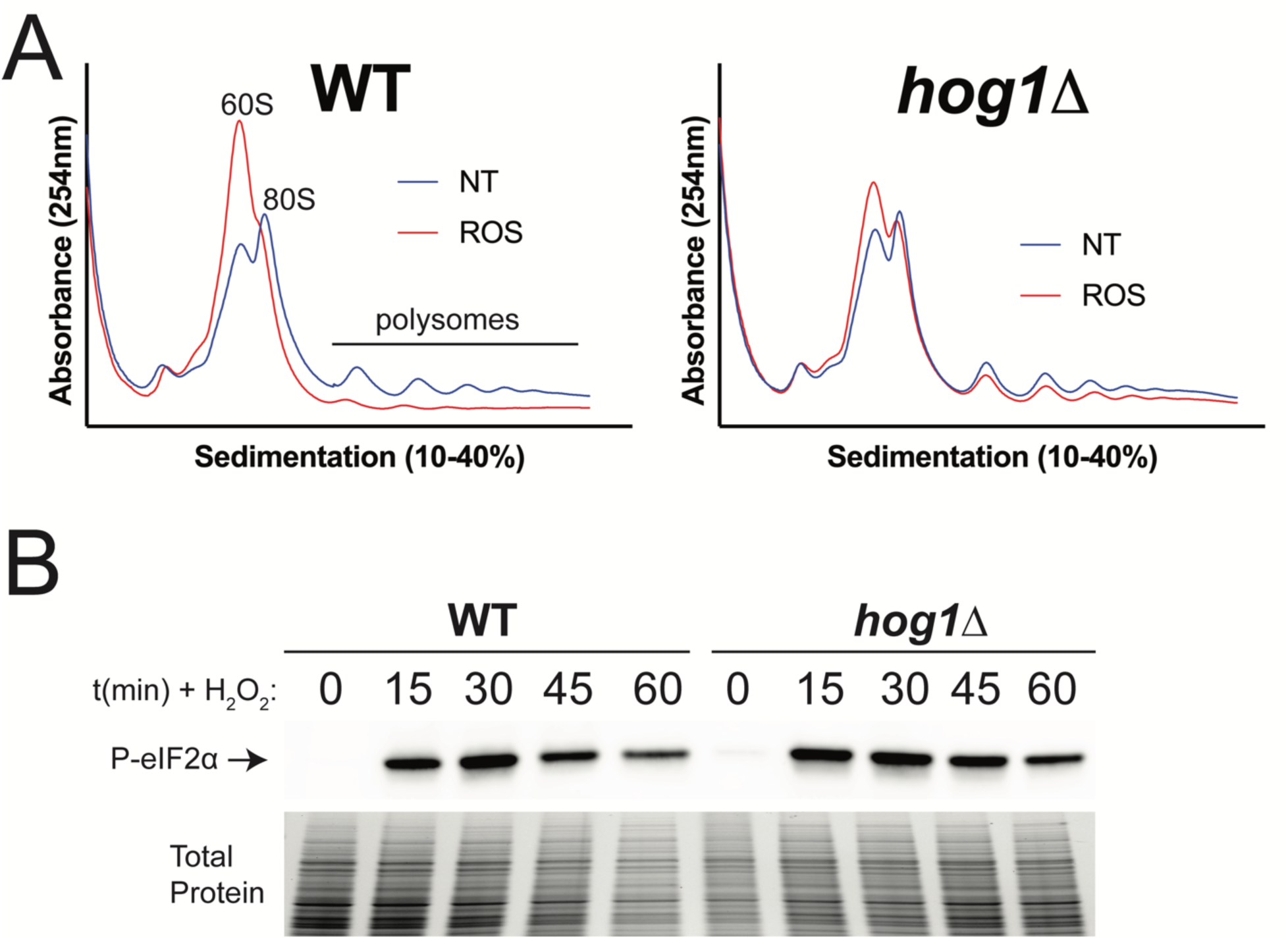
Loss of Hog1 results in defects in translation repression in response to ROS. WT and *hog1*Δ cells were grown to midlogarithmic phase, followed by addition of 2mM H_2_O_2_ to simulate oxidative stress. (A) RNA extracts were prepared from treated and untreated cells, followed by analysis of translation by polysome profiling. NT = no treatment, ROS = 30 mins of treatment with 2mM H_2_O_2_. (B) Protein extracts were prepared from cells at the indicated time points and probed using an antibody against phospho-eIF2α. Total protein is shown as a loading control (*n = 3*).

To ask whether altered translation repression in *hog1*Δ could be attributable to changes in Gcn2 output, we next investigated whether P-eIF2α in response to ROS was altered in *hog1*Δ. To study this, we probed protein lysates from WT and *hog1*Δ strains treated with 2mM H_2_O_2_ for P-eIF2α by western blotting (**Fig. 2B**). We found that both strains robustly phosphorylated eIF2α in response to oxidative stress, and saw no appreciable difference in the response of WT and *hog1*Δ. Together, these results suggest that reduced translation repression in *hog1*Δ does not result from altered eIF2α phosphorylation.

### RNA-sequencing of cells treated with 2mM H_2_O_2_ reveals large-scale reprogramming of the transcriptome

To better understand the genetic interaction between Hog1 and Gcn2 during oxidative stress, we harvested RNA from WT, *hog1*Δ, *gcn2*Δ, and *hog1*Δ*gcn2*Δ strains in the presence or absence of 2mM H_2_O_2_ and performed poly(A)-purified RNA sequencing on an Illumina platform. Principal component analysis (PCA) was performed on the resulting data, and indicated that much of the variability was due to treatment with 2mM peroxide and deletion of Hog1 (**Fig. S1**). The first two principal components appeared to account for these differences, which accounted for >80% of variability in our data (**Fig. S1A-B**). While deletion of Gcn2 appeared to have a smaller effect on variability along these principal components than Hog1 deletion, differences between WT and *gcn2*Δ were clearly differentiable along other principal components (**Fig. S1C**).

Differential expression was calculated in a pair-wise manner between samples, using a fold-change threshold of 1.75 and adjusted p-value < 0.05 as a significance threshold. We found that the response of WT cells was very robust, with over half of the transcriptome undergoing significant changes in expression in response to 2mM H_2_O_2_ (**Fig. 3A, S2A**). While this response appeared evenly distributed between upregulation and downregulation of transcripts, there was a greater magnitude of change among upregulated transcripts. Consistent with the role of Hog1 and Gcn2 as regulators of stress responses, the vast majority of differentially expressed transcripts in our mutants, relative to WT, were found only under the treated conditions (**Fig. S2B**). In light of this, we focused our analyses on transcripts that were dysregulated under ROS-treated conditions.

**Figure 3:**
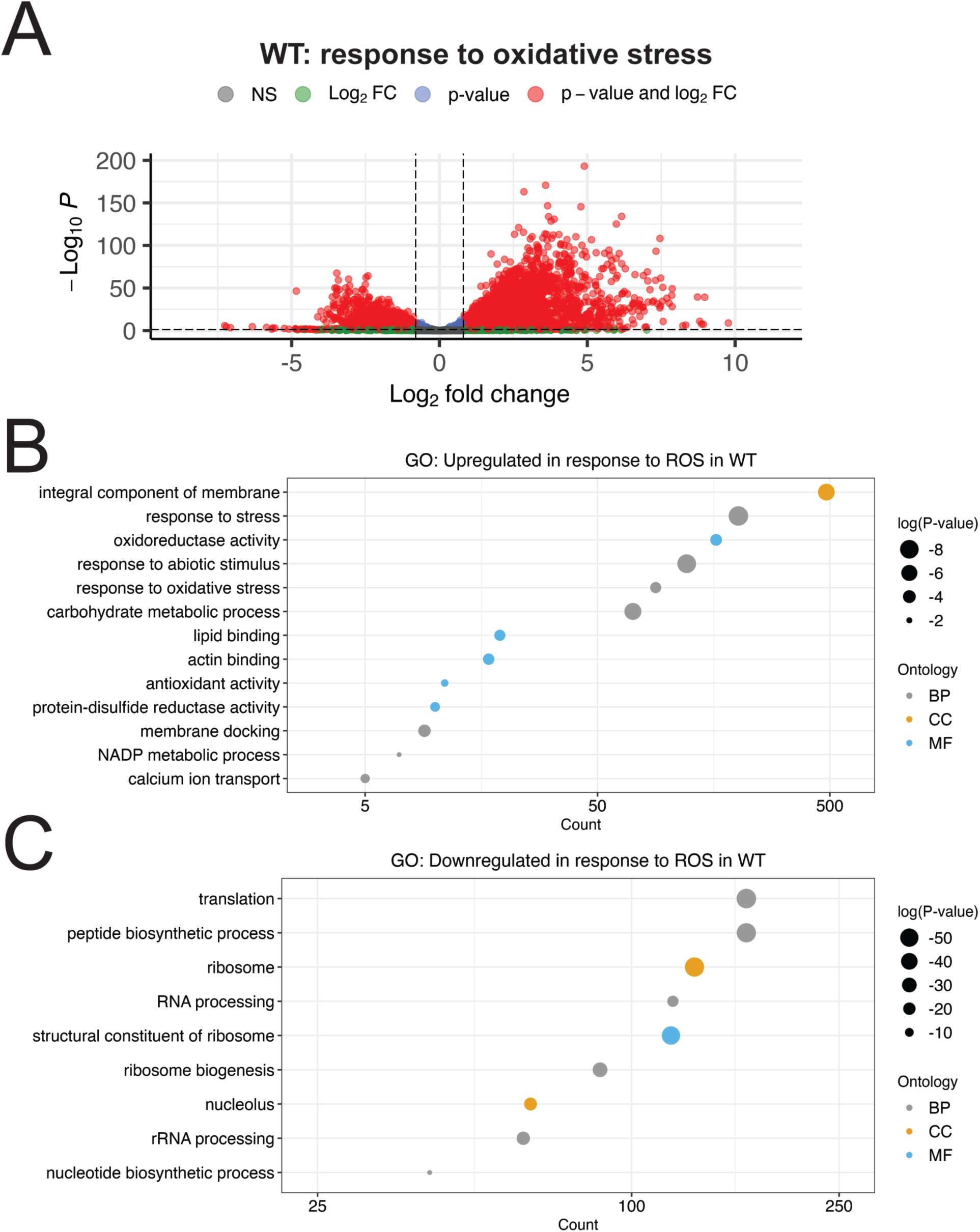
Overview of the WT response to ROS. (A) Volcano plot showing differential expression between WT cells treated with 2mM H_2_O_2_ and untreated cells. Differentially expressed transcripts were defined by abs(log2FC) > 0.81 and p-adj < 0.05, and are highlighted in red. (B-C) GO Analysis was performed on the set of transcripts that were (B) upregulated, or (C) downregulated in the WT response to ROS. The number of transcripts in each set belonging to that GO term is indicated along the x-axis, which is log-scaled. The size of each point is proportional to the significance value for enrichment, while the color of the point indicates the ontology from which that term is derived; BP = biological process, CC = cellular component, MF = molecular function.

We performed gene ontology (GO) enrichment analysis on genes upregulated by ROS treatment in the WT background, and found that genes related to the cellular response to stress were substantially enriched, including those encoding oxidoreductases and other antioxidants (**Fig. 3B, Table S1**). Also enriched among these genes were those encoding membrane-associated proteins and lipid binding, as well as regulators of metabolism, suggesting that morphological and metabolic reprogramming is likely involved in the response to 2mM H_2_O_2_.

Among transcripts downregulated by WT in response to oxidative stress, we observed a strong enrichment for proteins with ribosomal and translation-associated functions, as well as nucleic acid processing functions (**Fig. 3C**). This included ribosomal protein (RP) transcripts (**Table 1**), which are abundant under unstressed conditions, and need to be downregulated for ribosomal reallocation to stress response transcripts (11).

**Table 1:** Differential expression of ribosomal protein transcripts during the oxidative stress response.

| Gene | Change in Expression during ROS response (log<sub>2</sub>FC) |  |  |  |  |
| --- | --- | --- | --- | --- | --- |
|  | WT | *hog1Δ* | *gcn2Δ* | *hog1Δgcn2Δ* #9 | *hog1Δgcn2Δ* #11 |
| *RPS4* | -2.23 | -1.12 | -0.92 | -0.07 | 0.05 |
| *RPL7* | -2.22 | -1.12 | -1.03 | -0.14 | -0.05 |
| *RPS6* | -2.28 | -1.20 | -1.76 | -0.11 | -0.02 |
| *RPL3* | -2.37 | -1.48 | -1.74 | -0.31 | -0.25 |
| *RPS9* | -2.03 | -1.01 | -0.53 | 0.01 | 0.19 |
| *RPS1* | -2.22 | -1.07 | -0.89 | 0.01 | 0.06 |
| *RPL4* | -2.14 | -1.07 | -1.43 | -0.21 | -0.12 |
| *RPL8* | -2.24 | -1.23 | -1.18 | -0.13 | -0.02 |
| *RPL13* | -2.43 | -1.16 | -1.52 | -0.16 | -0.07 |
| *RPS2* | -2.42 | -1.18 | -1.05 | -0.14 | 0.05 |

Transcripts under the GO term "structural constituent of the ribosome" were selected, and the ten most highly expressed in unstressed WT were selected based on our RNA-seq data. The log<sub>2</sub>(fold-change) was calculated for each strain using DEseq2, comparing ROS-treated to unstressed cells.

### Gcn2 deletion is associated with reduced levels of transcripts encoding stress response regulators and amino acid biosynthesis transcripts

While PCA of our samples suggested that the WT and *gcn2*Δ strains behaved relatively similarly in response to oxidative stress (**Fig. S1**), we were nonetheless interested in understanding whether transcriptomic differences could help to explain the role of Gcn2 in the response to ROS. To do this, we first compared genes that were differentially expressed in the WT response to ROS to those differentially expressed in the *gcn2*Δ response to ROS, accounting for the directionality of the change in their expression in each background.

We observed a high degree of overlap in the response of WT and *gcn2*Δ strains (**Fig. S3**). Among those transcripts upregulated in response to oxidative stress in WT cells, more than 80% of transcripts were also upregulated in *gcn2*Δ (**Fig. S3A**). We performed GO analysis on the subset of transcripts which were induced by WT, but not *gcn2*Δ, and found that genes involved in virulence factor production and stress response were enriched (**Fig. S3B**). Among transcripts downregulated in response to oxidative stress in WT cells, ∼70% were also downregulated in *gcn2*Δ (**Fig. S3C**). GO analysis of transcripts repressed in the WT response, but not in *gcn2*Δ, indicated that many of these targets are involved in translation and mitochondrial metabolism (**Fig. S3D**). Overall, this indicates that although the response of WT and *gcn2*Δ to oxidative stress is similar, there are defects in induction of stress-related transcripts and repression of pro-growth mRNA in *gcn2*Δ that may underlie its sensitivity to oxidative stress.

Although genes induced by peroxide treatment in WT and *gcn2*Δ were mostly overlapping, we were also interested in defining whether these responses differed in extent, which could be accomplished through direct comparison of transcript levels in ROS-treated WT versus ROS-treated *gcn2*Δ.

Interestingly, we found that ∼2100 transcripts, nearly 25% of the transcriptome, met our cutoffs for differential expression between these two samples. GO analysis of transcripts that had lower expression in *gcn2*Δ compared to WT during oxidative stress showed an enrichment of transcripts involved amino acid biosynthesis, likely pointing to the ISR, and stress responses (**Fig. 4A**). GO analysis of transcripts that had greater expression in *gcn2*Δ compared to WT during oxidative stress indicated enrichment for genes with ribosome-or translation-associated functions, as well as protein folding (**Fig. 4B**). We found the latter finding intriguing, and noted that several chaperones had increased expression in *gcn2*Δ during oxidative stress, compared to WT (**Table S2**). We suspect that this could result from the inability to repress translation in the context of extensive damage, which could result in exposure of hydrophobic protein domains and induction of the heat shock response.

**Figure 4:**
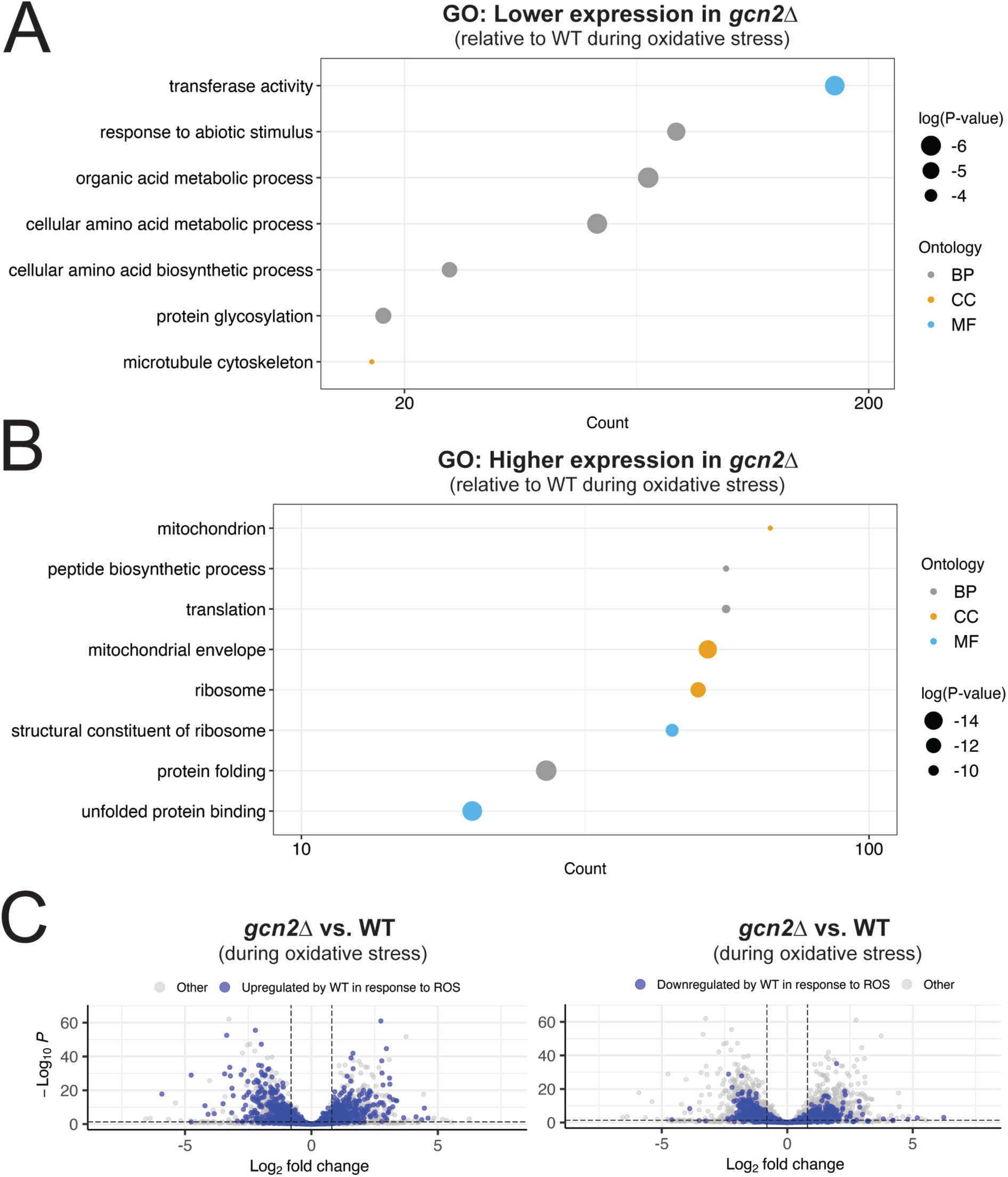
ISR induction and repression of translation-related mRNAs are defective in the absence of Gcn2. (A-B) GO analysis was performed on transcripts differentially expressed between ROS-treated *gcn2*Δ and ROS-treated WT. These transcripts were subsetted based on whether they were upregulated or downregulated in *gcn2*Δ with respect to WT, and visualized as described in Figure 3. (C) Differential expression between ROS-treated *gcn2*Δ and ROS-treated WT was visualized by volcano plot. Transcripts that were upregulated (left) or downregulated (right) in the WT response to ROS are highlighted in blue. Dashed lines represent thresholds for differential expression as described in the methods.

We next wanted to establish whether there was any global trend in dysregulation of the oxidative stress response in *gcn2*Δ compared to WT. To do so, we highlighted transcripts that were differentially expressed during the WT response to ROS in a volcano plot comparing *gcn2*Δ to WT during oxidative stress (**Fig. 4C**). For transcripts that were upregulated (**Fig. 4C**, left) or downregulated (**Fig. 4C**, right) in WT, while there was evidence of dysregulation in *gcn2*Δ, there was no clear trend in the data. This suggests that while there are differences in the *gcn2*Δ response to ROS, there was no global pattern indicative of a general defect in transcriptional induction or mRNA repression.

Lastly, since we previously defined the Gcn2-dependent ISR under oxidative stress using 1mM H_2_O_2_ (13), we were interested in looking at the induction of this response in the present study, which was performed using a higher concentration of hydrogen peroxide. To ask this, we visualized the behavior of ISR-dependent transcripts in the WT response by volcano plot (**Fig. S4**). Surprisingly, we did not observe robust induction of the ISR under 2mM H_2_O_2_ stress in WT cells. Given the extent of translation repression observed in WT cells at this higher concentration of peroxide (**Fig. 2A**), this result may indicate that there is an optimal extent of translation repression for ISR induction, and our experimental conditions exceeded this threshold. Alternatively, since we observed widespread transcript downregulation in the response to 2mM H_2_O_2_ (**Fig. 3A**), this result could indicate that ISR targets were destabilized during the oxidative stress response, masking the effect of ISR induction. The latter possibility is supported by direct comparison of *gcn2*Δ and WT during oxidative stress, which indicates that Gcn2 still has a positive effect on ISR transcript levels under these conditions (**Fig. 6C**, top right).

Overall, Gcn2 deletion appears to be associated with dysregulation of several classes of transcripts. Most notably, these include defects in induction of transcripts encoding effectors in stress responses, protein folding, and the ISR, as well as defects in repression of ribosomal protein transcripts.

### Hog1 deletion is associated with defects in the general response to ROS

To begin our analysis of the Hog1 deletion mutant, we first compared transcripts differentially expressed in the WT response to ROS to those differentially expressed in the *hog1*Δ response to ROS (**Fig. S5**). We found that ∼50% of genes upregulated in the WT background were also upregulated in *hog1*Δ in response to 2mM H_2_O_2_ (**Fig. S5A**). Among those transcripts which *hog1*Δ failed to induce, GO analysis indicated enrichment for functions in stress responses and protein post-translational modification (PTM), including several components the glutathione system (**Fig. S5B, Table S3**). Among transcripts downregulated in the WT background, less than 50% were downregulated in *hog1*Δ (**Fig. S5C**). GO analysis of transcripts which *hog1*Δ failed to repress indicated that these were enriched for functions in the ribosome, translation, and amino acid biosynthesis (**Fig. S5D**). Collectively, these data indicate that the response of WT and *hog1*Δ are dissimilar, and the sensitivity of *hog1*Δ to ROS could be dependent on dysregulation of the stress response, antioxidant systems, and repression of basally expressed transcripts.

We next set out to evaluate whether these responses differed in extent, again by direct comparison of ROS-treated WT and ROS-treated *hog1*Δ. We found that ∼3300 transcripts, nearly 40% of the transcriptome were differentially expressed between these samples. GO analysis of transcripts with lower expression in *hog1*Δ compared to WT during oxidative stress were enriched for regulators of stress responses, antioxidants, catalases, and calcium transport, as well as carbohydrate catabolism, which is a canonical target of the Hog1 pathway (**Fig. 5A**). Differential expression of antioxidants in WT versus *hog1*Δ during oxidative stress are summarized in **Table S1**. GO analysis of transcripts that were upregulated in *hog1*Δ compared to WT during oxidative stress were enriched for functions in the ribosome, translation, and amino acid biosynthesis (**Fig. 5B**). These results indicate that Hog1 plays an important role in induction of ROS-mitigating transcripts such as catalases, antioxidants, and other stress response factors, as well as repression of pro-growth mRNAs.

**Figure 5:**
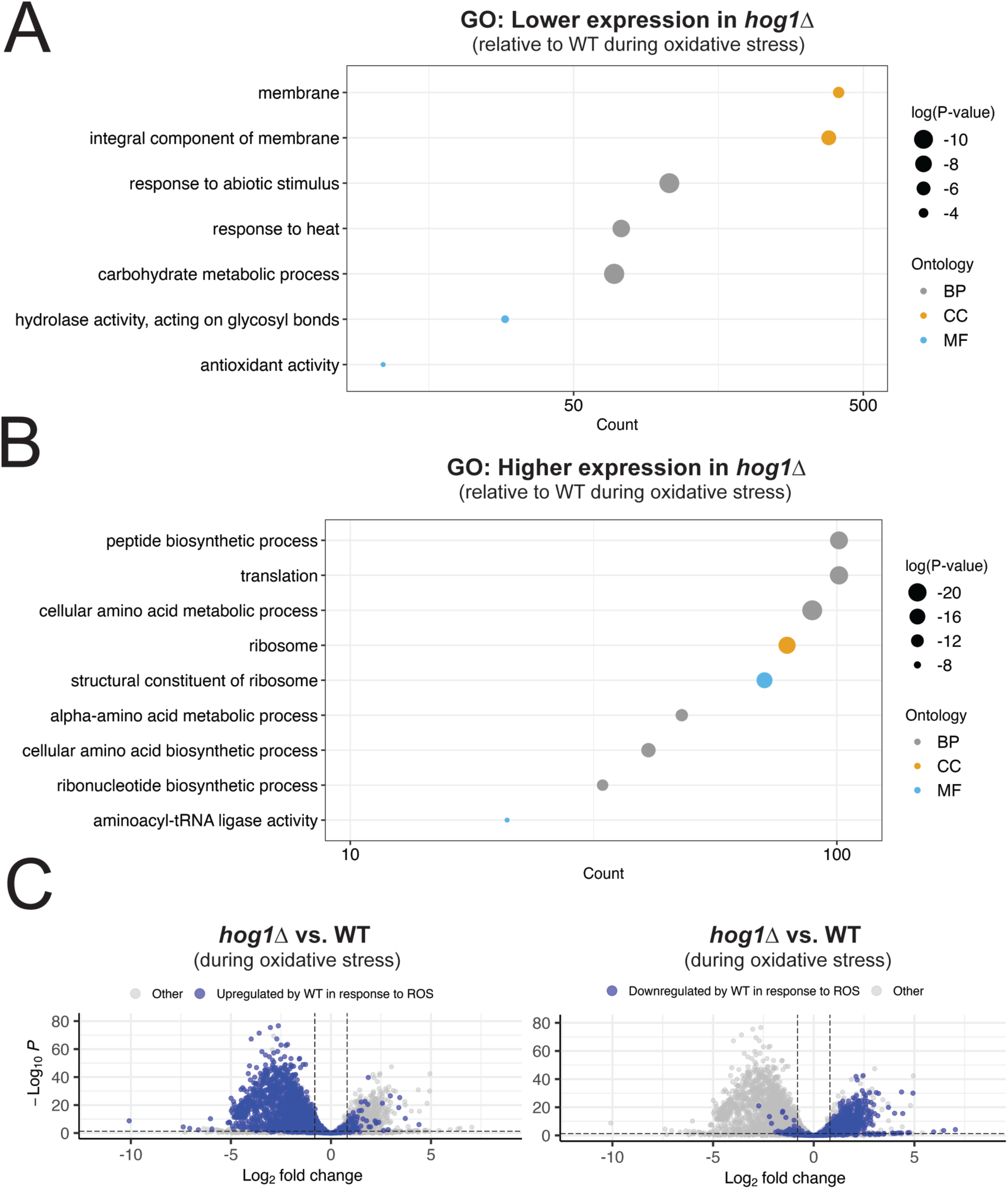
Stress response induction and transcript repression are globally dysregulated in the absence of Hog1. (A-B) GO analysis was performed on transcripts differentially expressed between ROS-treated *hog1*Δ and ROS-treated WT. These transcripts were subsetted based on whether they were upregulated or downregulated in *hog1*Δ with respect to WT, and visualized as described in Figure 3. (C) Differential expression between ROS-treated *hog1*Δ and ROS-treated WT was visualized by volcano plot. Transcripts that were upregulated (left) or downregulated (right) in the WT response to ROS are highlighted in blue. Dashed lines represent thresholds for differential expression as described in the methods.

**Figure 6:**
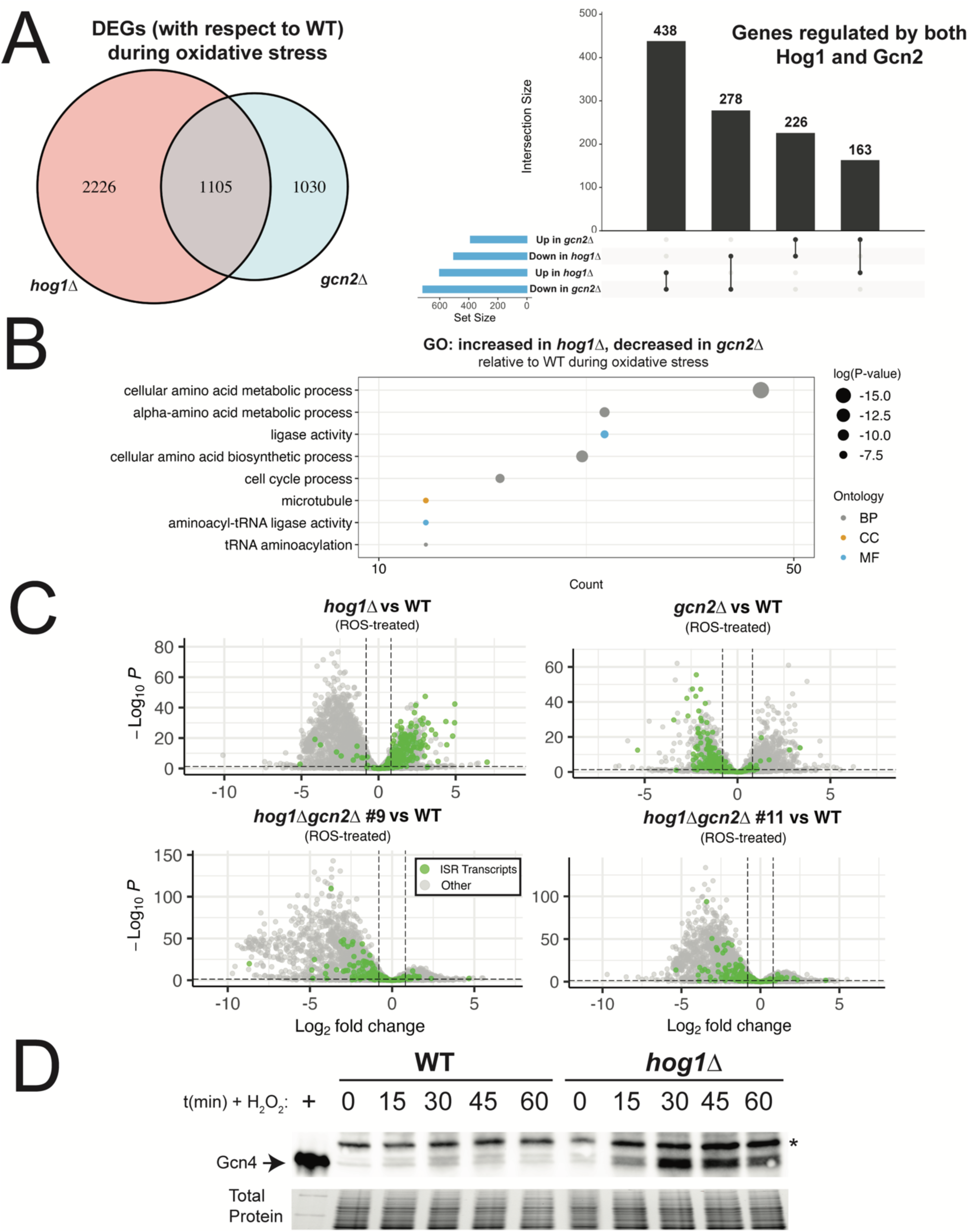
Hog1 deletion results in Gcn2-dependent upregulation of the ISR during oxidative stress. (A, left) Venn diagram comparing transcripts differentially expressed in ROS-treated *hog1*Δ or *gcn2*Δ, with respect to WT. (A, right) Upset plot visualizes the intersections between Hog1-and Gcn2-dependent transcripts, based on whether they are upregulated or downregulated. (B) GO analysis of transcripts that were upregulated in *hog1*Δ vs WT (during oxidative stress) and downregulated in *gcn2*Δ vs WT (during oxidative stress) was visualized as described in Figure 3. (C) Volcano plot of differential expression between each mutant and WT upon ROS treatment. Transcripts identified as targets of the ISR in a previous analysis are highlighted in green. Dashed lines represent thresholds for differential expression as described in the methods. (D) WT and *hog1*Δ were grown to midlogarithmic phase, followed by treatment with 2mM H_2_O_2_ for the indicated amounts of time. Protein lysates were probed by western blot using an anti-Gcn4 antibody. * = non-specific band. Total protein is shown as a loading control (*n = 3*).

Next, we were interested in determining whether there were global trends in dysregulation of the ROS response as a result of Hog1 deletion. To do this, we highlighted transcripts that were differentially expressed in the WT response to ROS in a volcano plot comparing *hog1*Δ to WT during oxidative stress (**Fig. 5C**). Among genes that were upregulated in the WT response to 2mM H_2_O_2_, the vast majority of these genes had lower expression in *hog1*Δ compared to WT (**Fig. 5C**, left). Among genes that were downregulated in the WT response to 2mM H_2_O_2_, there was a clear trend toward higher expression in *hog1*Δ compared to WT (**Fig. 5C**, right). Together, this result suggests that there is global dysregulation of the response to ROS in *hog1*Δ, with dampened induction of ROS-responsive transcripts, as well as defective repression of pro-growth transcripts.

Overall, we found that Hog1 deletion is associated with global dampening of the oxidative stress response, including canonical regulators of the ROS response such as catalases and other antioxidants.

We also found that transcript repression was defective in this background, which is a key step in ribosomal reallocation to stress response transcripts (11).

### The integrated stress response (ISR) is upregulated in *hog1*Δ during oxidative stress

Having established that Hog1 deletion is associated with transcriptome-wide defects in the response to oxidative stress, we next began to look for genetic interactions, convergence, and other points of similarity between Hog1 and Gcn2 deletion. To begin this analysis, we started by collecting genes that were dysregulated in *hog1*Δ or *gcn2*Δ, compared to WT, during oxidative stress. Of these genes, ∼1100 were shared by both strains (**Fig. 6A**).

When we accounted for the directionality of dysregulation in each mutant, these transcripts could be broken into four categories (**Fig. 6A**). Transcripts with increased expression in both *hog1*Δ and *gcn2*Δ during oxidative stress were greatly enriched for components of the ribosome, translation, and mitochondrial function (**Fig. S6A**), as would be expected based on the similarity of individual GO analyses for these strains (**Fig. S3D, S5D**). Among transcripts that had decreased expression in both mutants, there was enrichment of stress response regulators, although many of these were of unknown function or did not direct us toward a particular pathway (**Fig. S6B, Table S4**). Notably, the mRNA encoding thioredoxin reductase, *TRR1*, was among these transcripts.

Interestingly, the majority of differentially expressed transcripts in *hog1*Δ and *gcn2*Δ were dysregulated in opposite directions. Among those with increased expression in *gcn2*Δ and decreased expression in *hog1*Δ, GO analyses did not identify functional enrichment that met statistical cutoffs for significance. Manual review of this subset of genes, however, found that putative oxidoreductases, mitochondrion-associated proteins, and detoxifying enzymes were among these genes (**Table S5**). The largest category, which contained 438 transcripts, had increased expression in *hog1*Δ and decreased expression in *gcn2*Δ (**Fig. 6A**). GO analysis of these transcripts indicated that amino acid biosynthetic pathways were robustly enriched in this set of genes (**Fig. 6B**). This indicated to us that the Gcn2-dependent ISR is upregulated in *hog1*Δ compared to WT. This result was surprising, given our previous observation that P-eIF2α in response to ROS is unchanged in *hog1*Δ (**Fig. 2B**).

Since we identified a list of ISR-regulated targets in a previous study (13), we next visualized the abundance of these transcripts in our mutants relative to WT upon peroxide treatment (**Fig. 6C**). We found that the ISR was robustly upregulated in *hog1*Δ compared to WT. As expected, the abundance of ISR transcripts was lower in *gcn2*Δ. Importantly, simultaneous deletion of Hog1 and Gcn2 prevented upregulation of these transcripts, suggesting that upregulation of the ISR in *hog1*Δ is still dependent on Gcn2.

In light of this surprising finding, we investigated whether increased ISR induction in *hog1*Δ was associated with increased production of the ISR transcription factor, Gcn4, in this background. To do this, we measured Gcn4 protein abundance by western blotting in WT and *hog1*Δ upon treatment of cells with 2mM H_2_O_2_ (**Fig. 6D**). Compared to WT, we observed higher levels of Gcn4 protein in *hog1*Δ upon oxidative stress. This result indicates that Hog1 is a regulator of ISR induction during oxidative stress.

Taken in conjunction with our observations in **Figure 2B**, this indicates that loss of Hog1 regulates induction of the ISR during oxidative stress without altering eIF2α phosphorylation.

Finally, since we observed dampened translation repression during oxidative stress in *hog1*Δ by polysome profiling (**Fig. 2A**), we next asked whether this could be permissive for increased translation of the *GCN4* mRNA in this background. To verify that persistent polysome complexes in *hog1*Δ were indicative of active translation, we measured translational output in WT and *hog1*Δ by puromycin incorporation assay (**Fig. S7**). Surprisingly, we observed comparable levels of translational output in WT and *hog1*Δ upon treatment with ROS. This suggested that persistent polysome complexes in *hog1*Δ are not necessarily indicative of active translation, and could represent mRNAs with stalled ribosomes. Thus, increased Gcn4 production in *hog1*Δ is unlikely to be the result of globally increased translation, and likely occurs via a different mechanism.

### The Hog1 and Gcn2 pathways converge on regulation of ribosome-related transcripts

Given the high degree of complexity involved in analyzing the effect of simultaneous deletion of Hog1 and Gcn2, we implemented a subsetting strategy to analyze differential expression in *hog1*Δ*gcn2*Δ (**Fig. 7A**). Since we sequenced two independent clones of the double knockout strain, we started by identifying transcripts that were differentially expressed in both double knockouts, with respect to WT, during oxidative stress. Then, we filtered from these lists any genes that were also dysregulated in *hog1*Δ or *gcn2*Δ, with respect to WT, during oxidative stress. The resulting list represents transcripts that were uniquely dysregulated in the double knockouts, but not either single knockout.

**Figure 7:**
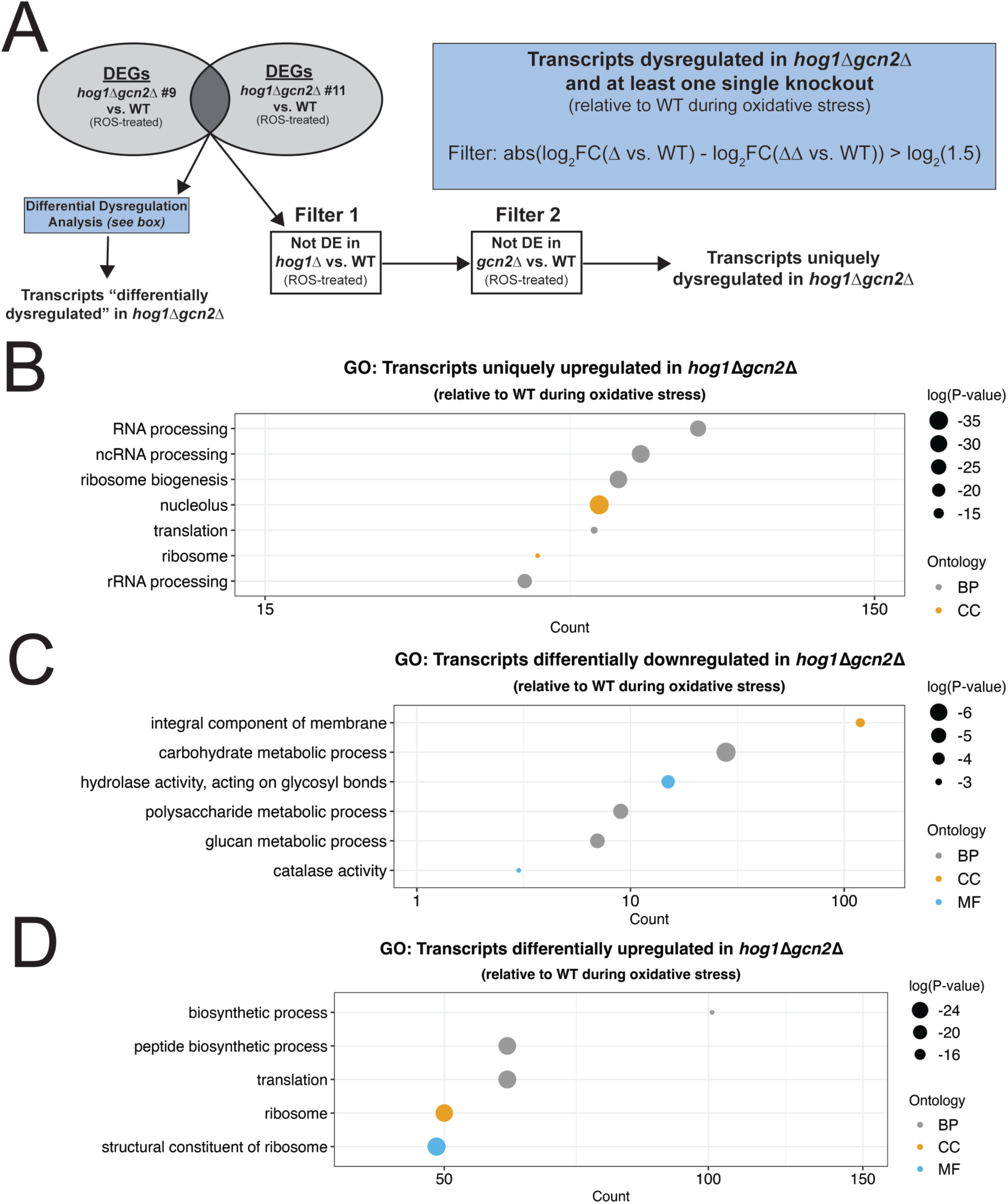
Loss of both Hog1 and Gcn2 results in emergent defects in repression of transcripts related to ribosome function. (A) Schematic showing filtering strategies for identification of transcripts with distinct patterns of dysregulation in *hog1*Δ*gcn2*Δ. (B) GO analysis was performed on transcripts that were uniquely upregulated in *hog1*Δ*gcn2*Δ compared to WT during ROS treatment and visualized as described in Figure 3. (C-D) Same as *(B)* except with transcripts that were differentially downregulated or differentially upregulated in *hog1*Δ*gcn2*Δ.

Among transcripts with uniquely lower expression in *hog1*Δ*gcn2*Δ, GO analysis yielded few results that were statistically significant. Manual review of these 314 transcripts found that transcripts involved in transport, metabolism, and regulation of the cytoskeleton were present (**Table S6**). GO analysis of the 640 transcripts with uniquely increased expression in *hog1*Δ*gcn2*Δ compared to WT, however, identified robust enrichment for genes involved in ribosome biogenesis and RNA processing (**Fig. 7B**). This suggests that loss of both Hog1 and Gcn2 produces an emergent defect in regulation of ribosome biogenesis and processing factors.

While the previous analysis identifies transcripts that are uniquely dysregulated in *hog1*Δ*gcn2*Δ with respect to WT, we also wanted to consider whether there were transcripts that differed in their extent of dysregulation in the double knockout, even if they were also dysregulated in single knockouts. To do this, we implemented a difference-of-differences-like approach. In short, from the subset of transcripts that were dysregulated in *hog1*Δ*gcn2*Δ and at least one of the single mutants, we selected transcripts where the extent of dysregulation in *hog1*Δ*gcn2*Δ was substantially different from what we observed in the single knockouts (**Fig. 7A**, see methods). Thus, this approach identified transcripts that were differentially dysregulated in *hog1*Δ*gcn2*Δ.

Among transcripts that were downregulated to a greater extent in *hog1*Δ*gcn2*Δ, we primarily saw enrichment for functions in carbohydrate metabolism, although three of the four catalases in *C. neoformans* were also present in this list (**Fig. 7C**). Among transcripts that were upregulated to a greater extent in *hog1*Δ*gcn2*Δ, we saw robust enrichment of transcripts with translation-and ribosome-associated functions (**Fig. 7D**). This suggested that while ribosome-related transcripts were dysregulated in *hog1*Δ and *gcn2*Δ, these transcripts were dysregulated to a unique extent in *hog1*Δ*gcn2*Δ, and likely targets of a genetic interaction between these two pathways.

Having identified several classes of transcripts where Hog1 and Gcn2 appear to play overlapping roles, we decided to visualize the behavior of some of these groups of genes by volcano plot (**Fig. 8**). The most ubiquitous trend in our GO analyses throughout this study was differential regulation of core components of the ribosome, suggesting that repression of these pro-growth transcripts was regulated simultaneously by Hog1 and Gcn2. We visualized the regulation of transcripts annotated under the GO term “structural constituent of the ribosome” by volcano plot (**Fig. 8A**). We found that while there was dampened repression of these mRNAs in *hog1*Δ and *gcn2*Δ during the ROS response, loss of both Hog1 and Gcn2 ablated repression of these transcripts entirely. To verify this result by another method, we performed northern blotting for the ribosomal protein transcript, *RPL2*, from RNA extracts from WT, *hog1*Δ, *gcn2*Δ, and *hog1*Δ*gcn2*Δ upon treatment with ROS (**Fig. 8B**). We saw no evidence of repression of *RPL2* over the course of one hour in *hog1*Δ*gcn2*Δ, whereas there was dampened repression in *hog1*Δ and *gcn2*Δ compared to WT, which agreed with our RNA-seq results.

**Figure 8:**
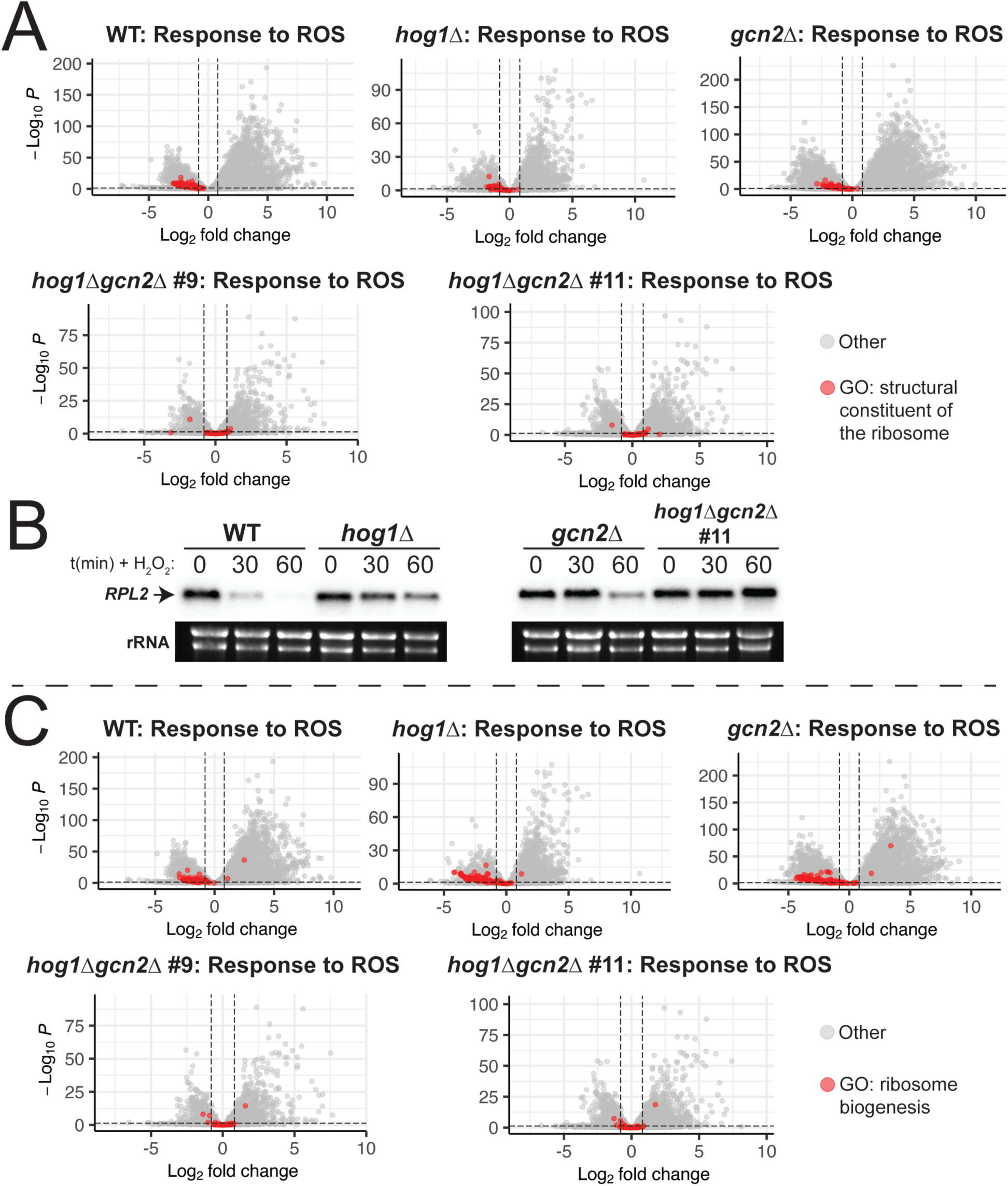
Visualization of transcripts with ribosome-related functions in *hog1*Δ*gcn2*Δ. (A,C) The response of each strain to 2mM H_2_O_2_ was visualized by volcano plot. All transcripts categorized under the indicated GO term are highlighted in red. Dashed lines represent thresholds for differential expression as described in the methods. (B) WT, *hog1*Δ, *gcn2*Δ, and *hog1*Δ*gcn2*Δ were grown to midlogarithmic phase and treated with 2mM H_2_O_2_ for the indicated amounts of time. RNA extracts were probed by northern blotting for the *RPL2* mRNA using the ^32^P labeled probe. rRNA is shown as a loading control (*n = 3*).

We next elected to visualize the behavior of transcripts encoding ribosome biogenesis factors, since this GO term was unique to our analysis of *hog1*Δ*gcn2*Δ (**Fig. 7B, 8C**). We found that transcripts with annotated functions in “ribosome biogenesis” were repressed in the WT background in response to oxidative stress. In *hog1*Δ and *gcn2*Δ, this repression appeared to be intact, whereas *hog1*Δ*gcn2*Δ did not appear to repress these transcripts at all. Overall, our results indicate that repression of transcripts with ribosome-associated functions are a major point of convergence between the Hog1 and Gcn2 pathways, as their downregulation is almost completely dependent on these pathways. Since repression of some of these transcripts still occurred in *hog1*Δ and *gcn2*Δ single mutants, our data suggests that there is a degree of functional redundancy between Hog1 and Gcn2 in promoting repression of these transcripts.

## Discussion

In this study, we examined the dependence of the *C. neoformans* oxidative stress response on the Hog1 and Gcn2 pathways. This led to a few important observations: First, we showed that loss of Hog1 and Gcn2 is highly consequential for the ability of *C. neoformans* to tolerate ROS. Second, we established that loss of Hog1 is associated with transcriptome-wide defects in mounting the response to ROS. Third, we demonstrated that induction of the ISR is regulated by Hog1, allowing for compensatory induction of this response in *hog1*Δ. Lastly, we found that the Hog1 and Gcn2 pathways are required for downregulation of transcripts encoding ribosome-associated factors.

Considered as a whole, our data indicate that Hog1 and Gcn2 are both global regulators of the response to ROS. In **Figure 1**, we demonstrated that the presence of at least one of these pathways is critical for ROS tolerance. Our RNA-seq data indicate that the WT response to these stressors is very robust, and involves not only upregulation of stress-related transcripts, but also downregulation of the unstressed growth program (**Fig. 3**). We measured dysregulation of this response in Hog1-and Gcn2-deleted backgrounds, and identified several functional classes of genes that are deficient in these mutants, including canonical regulators of ROS tolerance (**Fig. 4-5**). Notably, we also observed defects in repression of mRNAs encoding structural components of the ribosome (**Fig. 4-5, 8A-B**), which is a key step for ribosomal reallocation and energy conservation during translatome reprogramming (11). We also identified transcripts co-regulated by both of these pathways that are uniquely dysregulated in *hog1*Δ*gcn2*Δ, and found that these were most enriched for ribosome-associating factors (**Fig. 7B, 8C**).

These included not only structural components of the ribosome, but also factors involved in ribosome biogenesis and RNA processing, which did not appear to be dysregulated in *hog1*Δ or *gcn2*Δ alone.

A central goal of this study is to understand why *hog1*Δ*gcn2*Δ exhibits such a profound defect in ROS tolerance. Our observations in **Figure 5** indicate that the ROS response in *hog1*Δ is globally dysregulated, which we would expect to profoundly impact ROS tolerance. This prediction is disproven, however, by our observations in **Figure 1**, which showed that *hog1*Δ only exhibits modest sensitivity to oxidative stress. We believe this discrepancy is explained by our data in **Figure 6**, which noted a Gcn2-dependent upregulation of the ISR in *hog1*Δ. As we established in our previous study which defined the ISR regulon (13), many of the transcripts upregulated by this response are enriched for functions in amino acid biosynthesis and oxidoreductase activity. Additionally, many metabolites involved in amino acid biosynthesis regulate NADPH flux, supporting the idea that amino acid biosynthesis is closely linked to redox balance (28). Conversely, while we did not observe convincing evidence of compensatory Hog1 signaling in *gcn2*Δ, it is possible that Hog1 rescues this mutant due to its role as a general regulator of induction of ROS-mitigating transcripts.

While ISR induction in *hog1*Δ is one possible explanation for its ROS tolerance compared to *hog1*Δ*gcn2*Δ, we must also consider how other aspects of translatome reprogramming are dysregulated in this mutant. Our data indicate that loss of both Hog1 and Gcn2 results in profound defects in repression of transcripts that encode ribosomal components and biogenesis factors (**Fig. 7-8**). It has been shown previously in budding yeast that efficient translation of stress-related mRNAs requires removal of pro-growth mRNAs from the translating pool (11). In line with this, our lab previously reported that loss of the major deadenylase, Ccr4, which is required for repression of these mRNAs in *C. neoformans*, results in profound defects in stress tolerance and virulence (14, 29). Thus, it is possible that the inability to repress these transcripts, which are some of the most abundant in the cell, prevents *hog1*Δ*gcn2*Δ from mounting a successful stress response.

Another major question raised by this work is the mechanism underlying increased induction of the ISR in *hog1*Δ (**Fig. 6**). We observed increased levels of Gcn4 protein in *hog1*Δ during oxidative stress (**Fig. 6D**), despite normal levels of P-eIF2α in this mutant (**Fig. 2B**). Since increased ISR induction in *hog1*Δ was still dependent on the presence of Gcn2 (**Fig. 6C**), we suspect that Hog1 regulates another part of the Gcn2 pathway that only affects translation of the *GCN4* mRNA. Since our RNA-seq data did not provide evidence that the *GCN4* mRNA is differentially expressed in *hog1*Δ, we suspect that this could be result from changes to the stoichiometry of translation factors, which has been shown to alter ISR induction in other eukaryotes (30, 31). Indeed, we identified several translation factors with altered mRNA levels in *hog1*Δ, including subunits of eIF2B, the target of phosphorylated eIF2α (**Table S7**).

Additional study is warranted to better understand the genetic interaction between Hog1 and Gcn2. The present study, in conjunction with our previous study of Hog1-Gcn2 crosstalk during thermal stress (27), indicates that the nexus between these pathways is stress-specific. This implies that there are multiple mechanistically distinct interfaces between these pathways, which have biologically important consequences. Future studies are needed to define the feedback loops that exist between these pathways, and identify outputs of each pathway that are masked by functional redundancy. Additionally, given the exacerbated sensitivity of *hog1*Δ*gcn2*Δ to multiple host-derived stressors, we believe studies are warranted to evaluate the extent of virulence attenuation in this mutant.

Another important implication of this work is that it demonstrates the challenges associated with study of core conserved signal transduction pathways. We believe this work is of interest to anyone studying signal transduction because it demonstrates the extent to which homeostasis can be recalibrated when these core pathways are disrupted. Understanding these phenomena is necessary to properly assign which functions are attributable to which genes, and predict the effects of disrupting multiple pathways simultaneously. Given the urgent need for new antifungal strategies to treat cryptococcosis, we also believe these findings could prove useful in development of combined treatment strategies with greater efficacy.

## Materials and methods

### Strains and growth conditions

*C. neoformans* strain H99 was used as the wild-type background. The *hog1*Δ, *gcn2*Δ, and *hog1*Δ*gcn2*Δ mutants are derived from this background, as described previously (27). For all assays, cells were first grown overnight in YPD media (1% yeast extract, 2% peptone, and 2% dextrose; Difco) at 30°C with shaking at 250rpm in snap-cap tubes. For spot plate analyses, cells were used directly from this overnight culture. For all other assays, fresh YPD cultures were seeded at OD_600_ = 0.2, and grown to mid-logarithmic phase (OD_600_ = ∼ 0.65) in baffled flasks with shaking at 250rpm prior to treatment with peroxide. For peroxide treatments, H_2_O_2_ was added from a concentrated stock into actively growing cultures without replenishing the growth medium.

### Spot plate analysis

Overnight cultures of the indicated strains were washed in sterile water and resuspended to an OD_600_ = 1.0. A ten-fold serial dilution was performed, and 5μL of each dilution was plated on YPD agar (Difco), which was prepared with or without peroxide at the indicated concentration. Cells were photographed after incubation in a 30°C incubator for three days. Shown is one representative replicate (*n=3*).

### Polysome profiling

Cells of the indicated strains were grown to midlogarithmic phase, and control samples were collected. The remaining cells were treated with 2mM H_2_O_2_ for thirty minutes.

Samples were pelleted and flash-frozen in liquid nitrogen and stored at-80°C until processing. Cells were lysed by mechanical disruption in polysome lysis buffer (20 mM Tris-HCl pH 8.0, 140 mM KCl, 5 mM MgCl_2_, 1% Triton X-100, 0.5mg/mL heparin sodium sulfate, 0.1 mg/mL cycloheximide) and RNA concentration in lysate was measured on a NanoDrop One. 200μg RNA was loaded over a 10-40% sucrose gradient, followed by centrifugation at 39,000rpm for 2 hrs in a SW41-Ti swinging bucket rotor ultracentrifuge. Polysome profiles were collected by measuring absorbance at 260nm, as described previously (27). Shown is one representative replicate (*n = 3*).

### Puromycin incorporation

Puromycin incorporation was performed as described previously (32), followed by western blotting as described below using an anti-puromycin antibody (Millipore, MABE343) and HRP-conjugated anti-mouse secondary antibody (CST, #7076).

### Western blotting

Cells of the indicated strains were grown to midlogarithmic phase, and control samples were collected. The remaining cells were treated with 2mM H_2_O_2_ for the indicated amount of time, then pelleted and flash-frozen. Thawed cells were lysed, and SDS-PAGE was performed as described previously (27). P-eIF2α was probed using rabbit α-phospho-eIF2α (AbCam, ab32157) and an HRP-conjugated secondary antibody (CST, #7074). Gcn4 levels were detected with a custom-generated antibody (GenScript) raised in rabbits exposed to full-length recombinant protein. HRP signal was measured on a BioRad ChemiDoc MP using Clarity Max ECL substrate (Bio-Rad). Shown is one representative replicate (*n = 3*).

### Northern blotting

Cells of the indicated strains were grown to midlogarithmic phase, and control samples were collected. The remaining cells were treated with 2mM H_2_O_2_ for the indicated amount of time, then pelleted and flash-frozen. Thawed cells were lysed using the Qiagen RNeasy kit, following manufacturer instructions. 3μg of RNA from each sample was loaded into an RNA gel, and probed by northern blot against the *RPL2* transcript as previously described (27). rRNA was visualized as a loading control.

### RNA-sequencing

Cells of the indicated strains were grown to midlogarithmic phase, and control samples were collected. The remaining cells were treated with 2mM H_2_O_2_ for one hour, then pelleted and flash-frozen. RNA was extracted by mechanical disruption using glass beads in RLT buffer with 1% β-mercaptoethanol. Samples were then purified according using the Qiagen RNeasy kit, following manufacturer instructions including on-column DNase I digestion. rRNA integrity was visualized by gel electrophoresis, then samples were sent for library preparation, poly(A) purification, and RNA sequencing on an Illumina platform (Azenta). 350M paired-end reads were collected (2 x 150bp). Two biological replicates were sequenced for each sample. In **Fig S1-S2**, ROS-treated samples are indicated when the named is appended “_ROS”, while untreated control samples are referred to by the strain name alone.

### RNA-seq analysis

FASTQ files were processed to generate a counts matrix using the following software in command-line: Cutadapt/3.2 (adapter trimming and quality control), STAR/2.7.2b (alignment), and RSEM/1.2.20 (read counting) (33–35). The programs FastQC and multiQC were used along this pipeline for quality control and to monitor alignment. Cutadapt was run with quality filter of 10, and minimum read length of 1 to remove empty reads, and was specified to trim the Illumina universal adapter (AGATCGGAAGAG). STAR was run in read alignment mode with default parameters using the FungiDB H99 genome build (version 63) (36). RSEM was run with a forward probability of 0.5, again using the FungiDB H99 genome build (v63) (36). All subsequent analysis was performed in R.

For differential expression, read counts were rounded and analyzed in pair-wise fashion using DESeq2 (37). DESeq2 was used to output MA-plots and differential expression tables. The resulting differential expression tables were then filtered for an absolute value of log2(fold-change) of at least log_2_(1.75), and p-adjusted value < 0.05. Transcripts meeting these thresholds were considered to be differentially expressed for all analyses. Pairwise differential expression tables are provided in **S1** Appendix.

### Visualization of RNA-seq data

Visualization of RNA-seq data was performed in R (38).

*Principal component analysis (PCA).* Variance-stabilized counts were obtained using DESeq2, filtered for low expression, and the 2000 most variable transcripts were selected for analysis. PCA was performed using the prcomp command in R. The Scree plot was generated based on these results using the ggplot2 package in R (39), and indicated that the first five principal components were sufficient to account for 95% of variability in the data. These components were retained, and the PCA plots were generated for these principal components using ggplot2.

*MA plots, Venn diagrams, and Upset plots*. MA plots were generated using the DESeq2 package in R (37). Venn diagrams were generated using the VennDiagram package in R (40). Upset plots using the UpsetR package in R (41).

*Volcano plots*. Volcano plots for pairwise comparisons were generated using the R package EnhancedVolcano (42), with custom gene lists imported and colored as indicated. Transcripts colored in blue and described as up-or down-regulated by the WT ROS response were defined based on DEseq2 output for the comparison of WT treated with 2mM H_2_O_2_ versus untreated WT. The ISR target list was defined as described in a previous analysis (13). For volcano plots highlighting transcripts under specific GO terms, ontologies were obtained from FungiDB (v68) (36). The GO terms corresponding to “structural constituent of the ribosome” and “ribosome biogenesis” are stored under the identifiers GO:0003735 and GO:0042254. Dashed lines in volcano plots represent the significance and fold-change cutoffs described above for differential expression.

*GO analyses*. Bubble plots were generated using ggplot2 (39) based on GO analyses performed using the gene ontology tool on FungiDB (v68) (36). Point size represents the log10(p-value), and the x-axis shows counts and is log-scaled. Points were colored based on the ontology from which they originate. For tables, differential expression data were filtered based on: GO:0003735 (“structural constituent of the ribosome”, **Table 1**), GO:0016209 (“antioxidant activity”, **Table S1**), GO:0006457 (“protein folding”, **Table S2**), GO:0015305 (“protein-disulfide reductase activity”, **Table S3**), (GO:0033554 (“cellular response to stress”, **Table S4**). **Tables S5-S6** were manually curated, and the full set of genes is provided in **S2 Appendix**. **Table S7** was filtered based on the presence of “translation initiation factor” in the gene description provided by FungiDB (v68).

*Analysis of hog1Δgcn2Δ.* To identify transcripts that are uniquely dysregulated in *hog1*Δ*gcn2*Δ with respect to WT during oxidative stress, differentially expressed transcripts from each clone of ROS-treated *hog1*Δ*gcn2*Δ versus ROS-treated WT were intersected. Then, any transcripts which were dysregulated in ROS-treated *hog1*Δ or *gcn2*Δ versus ROS-treated WT were filtered out.

To identify transcripts that were “differentially dysregulated” in *hog1*Δ*gcn2*Δ, the same strategy as above was implemented, except those transcripts which were also dysregulated in *hog1*Δ and *gcn2*Δ were retained rather than filtered out. Then, the log2(fold-change) values of each ROS-treated mutant versus ROS-treated WT were compared. These results were filtered such that only transcripts where log2(fold-change) in both *hog1*Δ*gcn2*Δ clones was at least 1.5-fold different from each single knockout were retained, using the following formulas:

*abs(log2FC(hog1Δgcn2Δ_ROS vs. WT_ROS) - log2FC(hog1Δ_ROS vs. WT_ROS)) > log2(1.5) abs(log2FC(hog1Δgcn2Δ_ROS vs. WT_ROS) - log2FC(gcn2Δ_ROS vs. WT_ROS)) > log2(1.5)*

This analysis was repeated for data from each clone of *hog1*Δ*gcn2*Δ, and these results were intersected to increase stringency of this analysis. These results are referred to in the text as upregulated or downregulated “to a greater extent” in *hog1*Δ*gcn2*Δ.

## Acknowledgements

This work was funded by Institutes of Allergy and Infectious Disease at the National Institute of Health under award numbers R01AI131977 and F31AI169969. We would also like to acknowledge Azenta, which performed library preparation, poly(A) purification, and Illumina sequencing of our samples.

---

**Figure S1:**
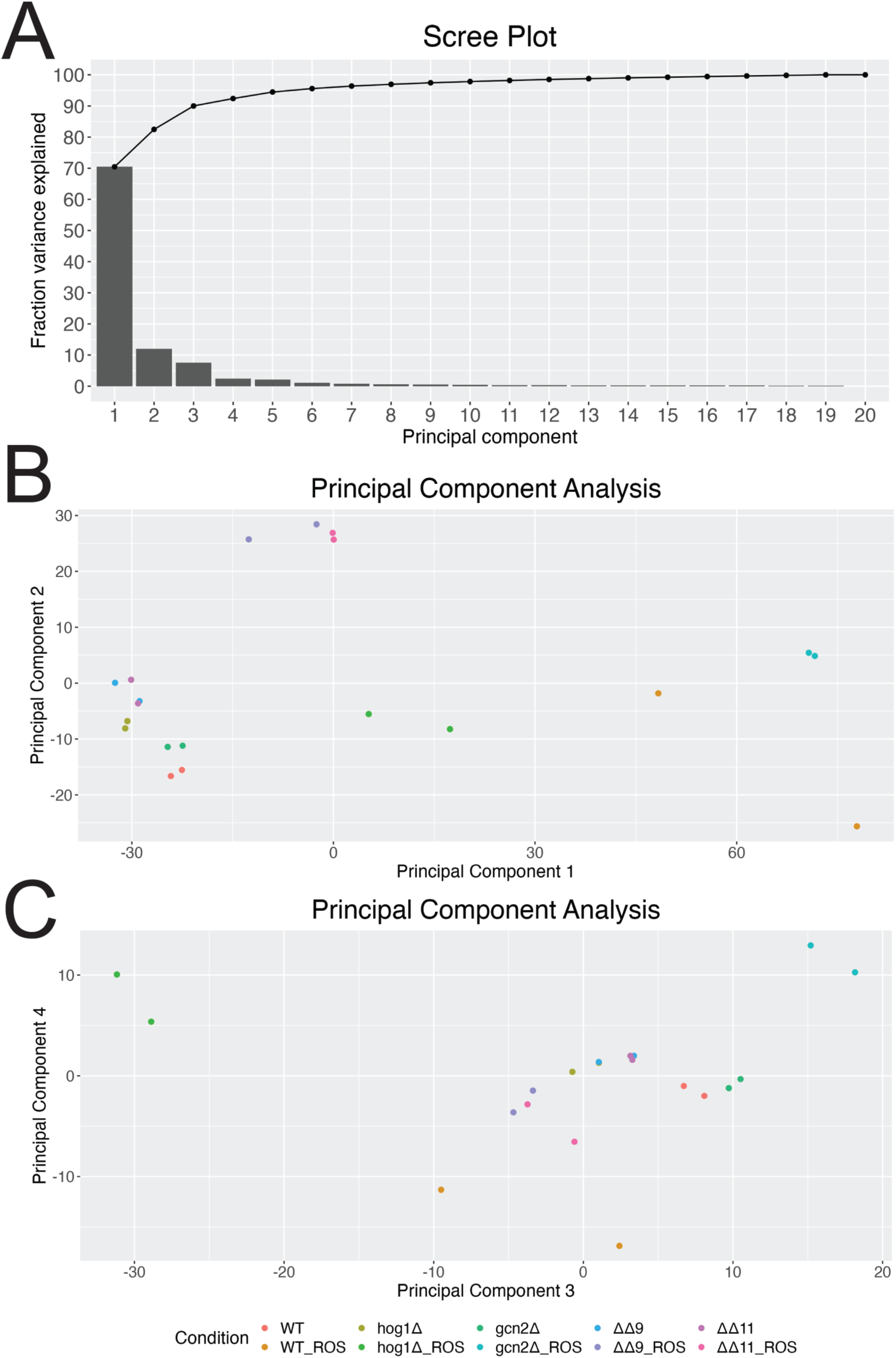
PCA of mRNA-seq data. PCA was performed on VST-transformed counts for the indicated samples using R. (A) Scree plot indicates the cumulative distribution by which principal components account for variability in the data. (B) PCA plot of Principal Components 1 and 2. (C) PCA plot of Principal Components 3 and 4. Samples with strain name alone are indicative of untreated control samples. Samples with names appended “_ROS” are indicative of cells treated with 2mM H_2_O_2_ for 1 hour.

**Figure S2:**
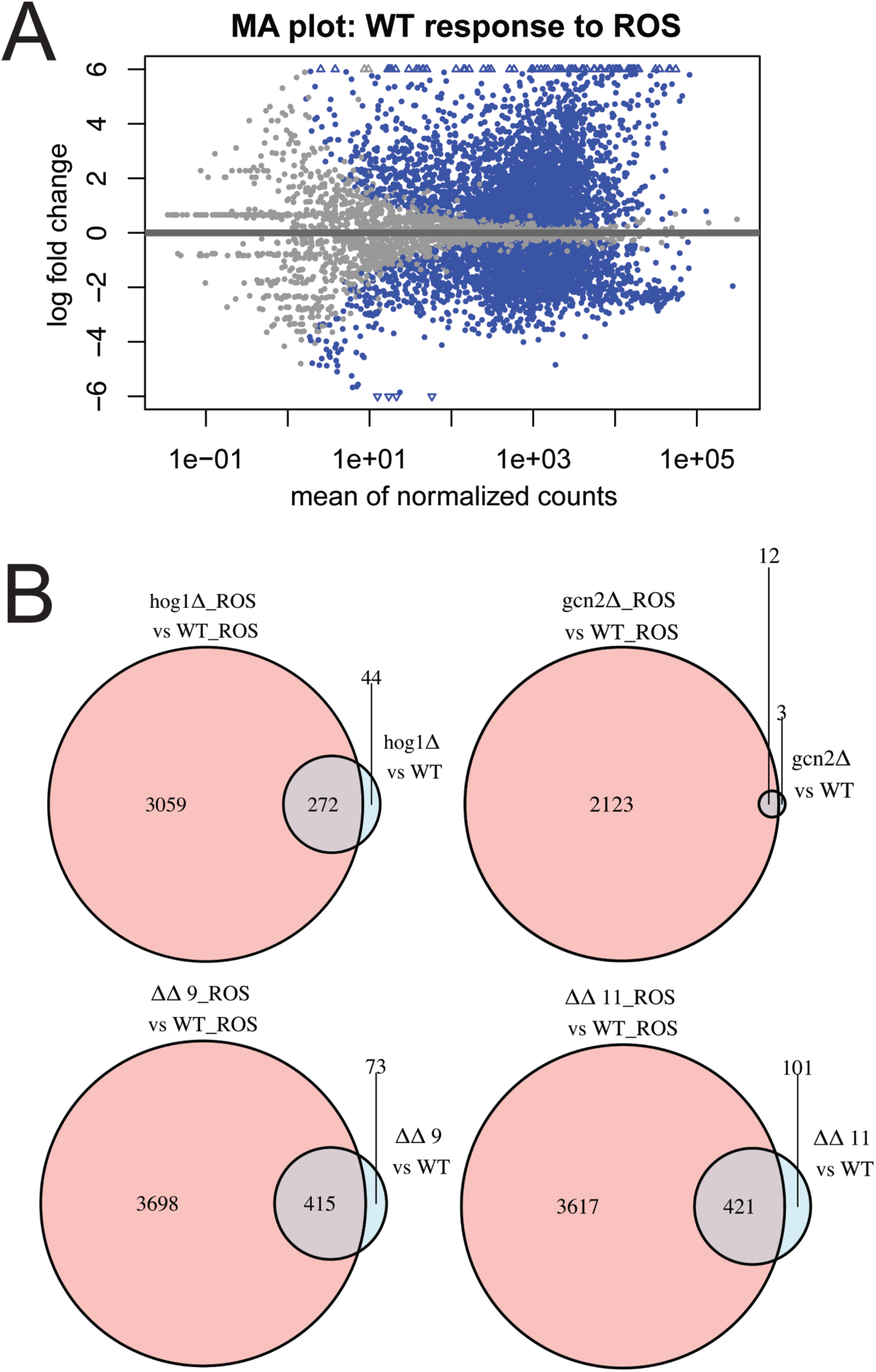
The *C. neoformans* ROS response involves robust changes to gene expression. (A) MA plot showing the WT response to ROS, with normalized counts on the x-axis and log2FoldChange on the y-axis. Blue dots represent significant changes to gene expression. (B) Summary of differential expression data between mutant strains and WT. Each set of Venn diagrams represents one strain, and compares the number of differentially expressed genes under untreated vs. treated conditions. Samples with names appended “_ROS” are indicative of cells treated with 2mM H_2_O_2_ for 1 hour.

**Figure S3:**
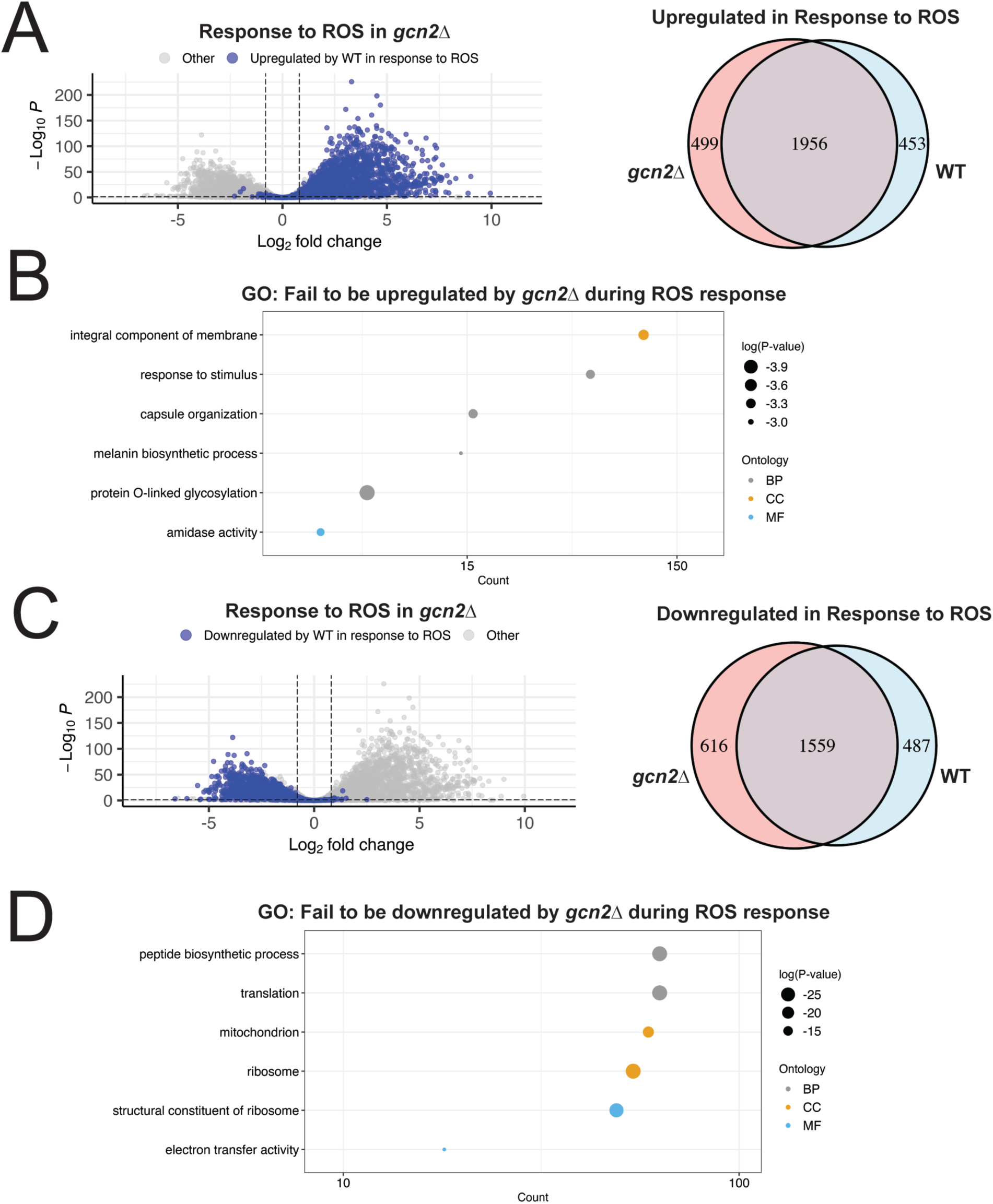
Comparison of the WT and *gcn2*Δ response to ROS. (A, left) Volcano plot showing the *gcn2*Δ response to treatment with 2mM H_2_O_2_, with genes upregulated in the WT response highlighted in blue. Dashed lines represent thresholds for differential expression as described in the methods. (A, right) Venn diagram comparing genes upregulated in the WT and *gcn2*Δ responses to ROS. (B) Transcripts that were unique to the WT response (i.e. transcripts that *gcn2*Δ failed to upregulate) were analyzed by GO and visualized as described in Figure 3. (C-D) The same analysis as in *(A-B)*, except focusing on transcripts that were downregulated in the stress response of each strain.

**Figure S4:**
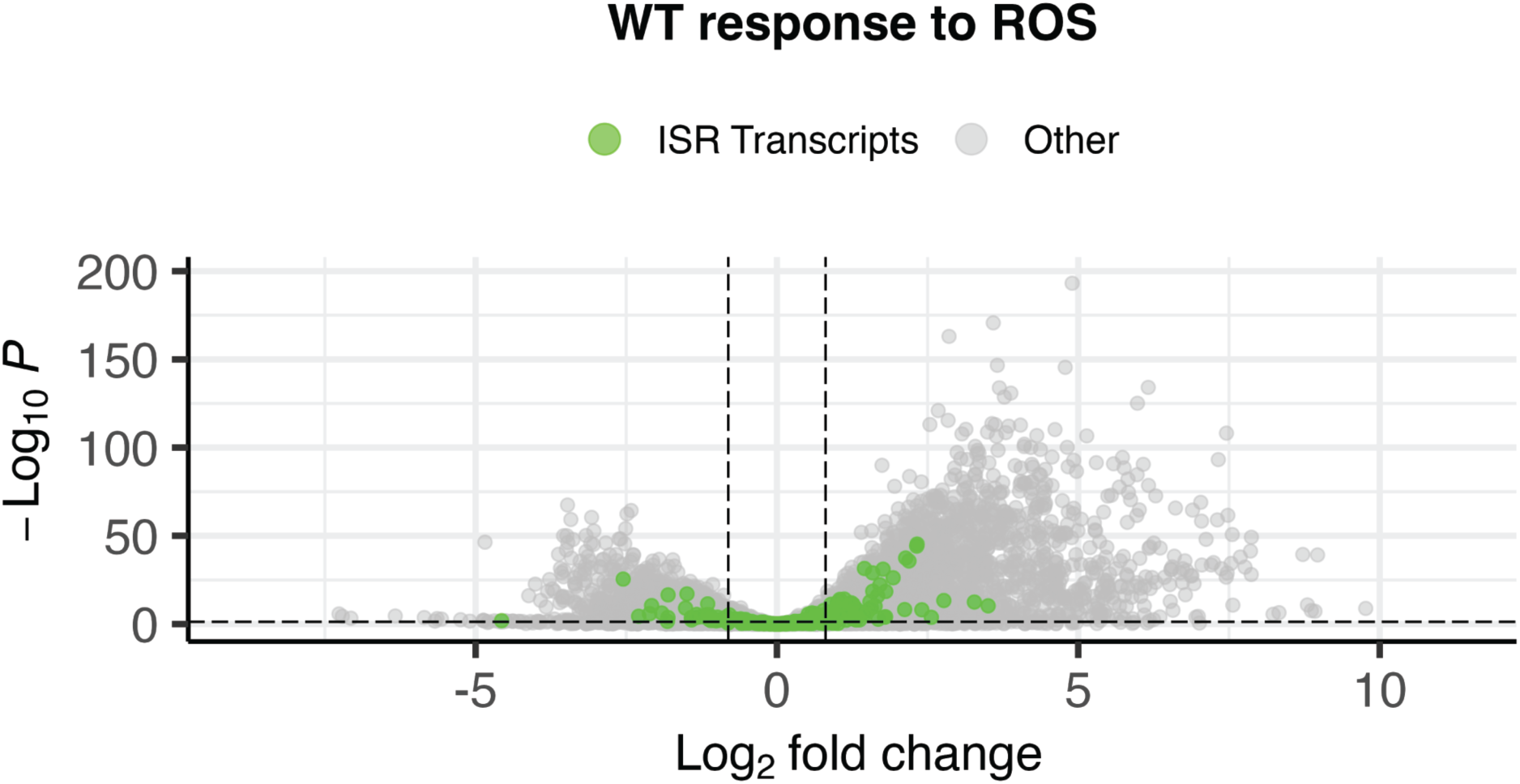
Visualization of the ISR in the WT response to 2mM H_2_O_2_. The WT response to treatment with 2mM H_2_O_2_ was visualized by volcano plot, with transcripts identified as targets of the ISR in a previous analysis highlighted in green. Dashed lines represent thresholds for differential expression as described in the methods.

**Figure S5:**
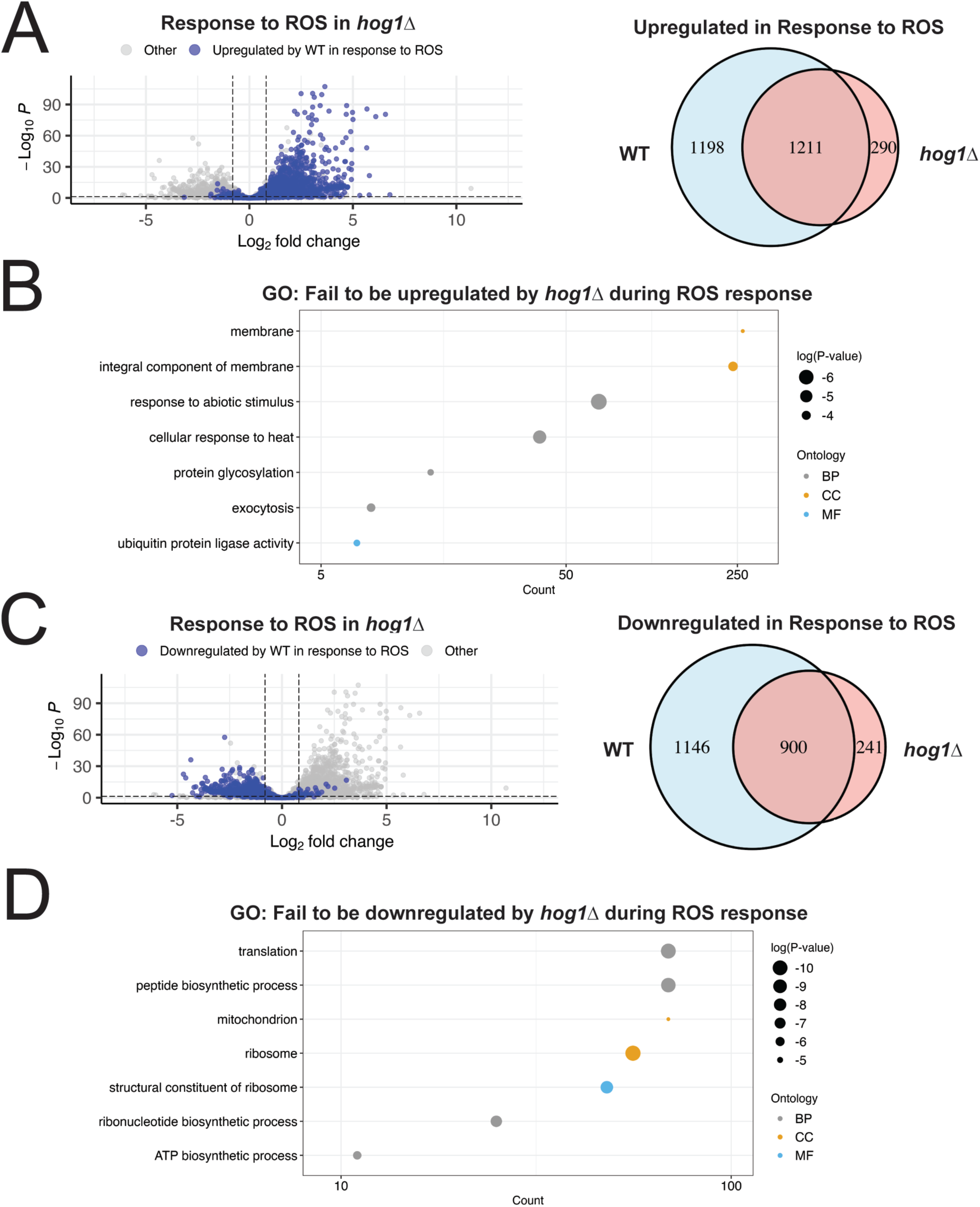
Comparison of the WT and *hog1*Δ response to ROS. (A, left) Volcano plot showing the *hog1*Δ response to treatment with 2mM H_2_O_2_, with genes upregulated in the WT response highlighted in blue. Dashed lines represent thresholds for differential expression as described in the methods. (A, right) Venn diagram comparing genes upregulated in the WT and *hog1*Δ responses to ROS. (B) Transcripts that were unique to the WT response (i.e. transcripts that *hog1*Δ failed to upregulate) were analyzed by GO and visualized as described in Figure 3. (C-D) The same analysis as in A-B, except focusing on transcripts that were downregulated in the stress response of each strain.

**Figure S6:**
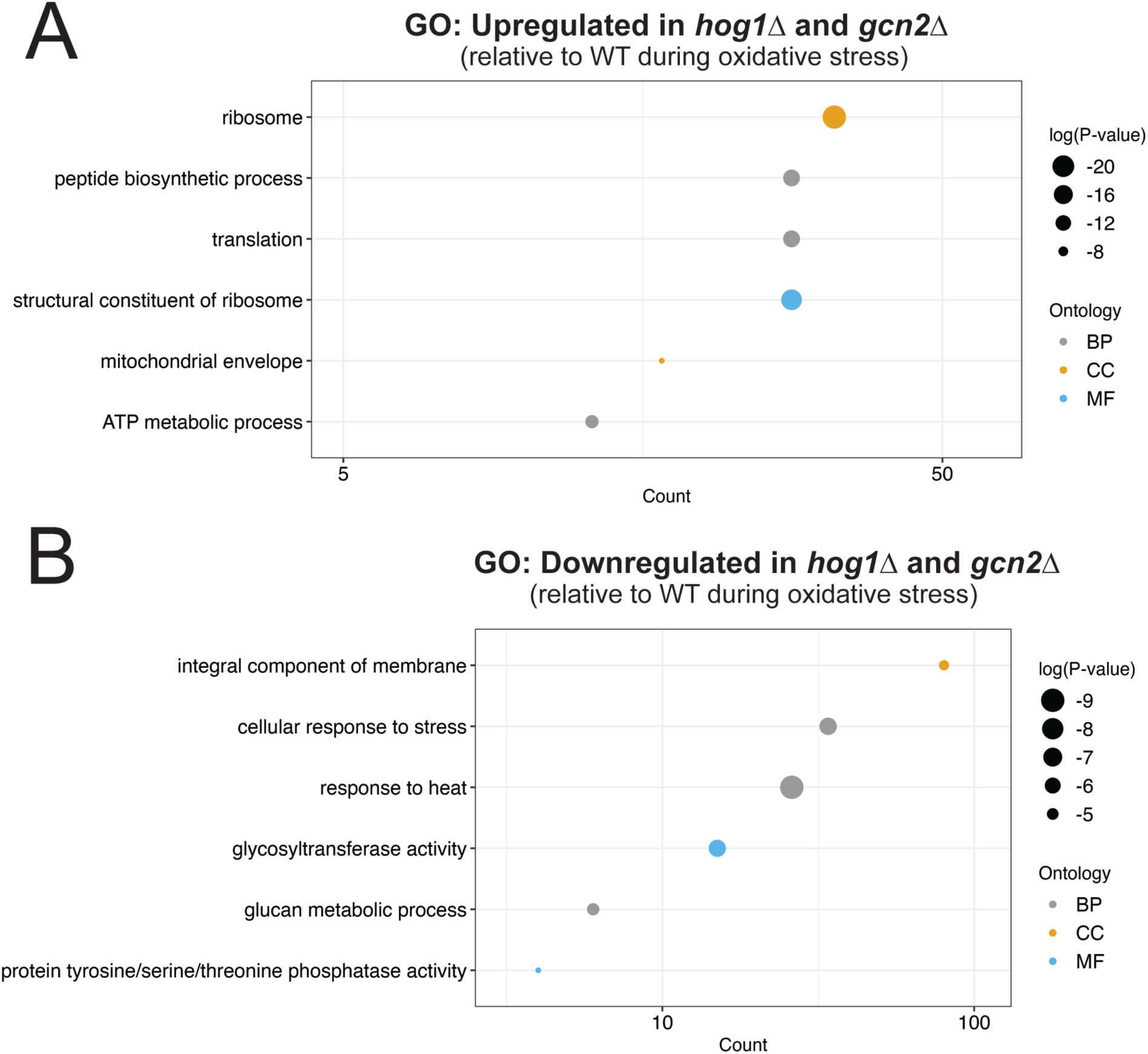
GO analysis of transcripts co-regulated by Hog1 and Gcn2 during ROS. (A-B) GO analysis was performed on transcripts differentially expressed in both *hog1*Δ and *gcn2*Δ, with respect to WT during ROS treatment. (A) GO analysis of transcripts upregulated in both *hog1*Δ and *gcn2*Δ, with respect to WT during ROS treatment was visualized as described in Figure 3. (B) GO analysis of transcripts downregulated in both *hog1*Δ and *gcn2*Δ, with respect to WT during ROS treatment was visualized as described in Figure 3.

**Figure S7:**
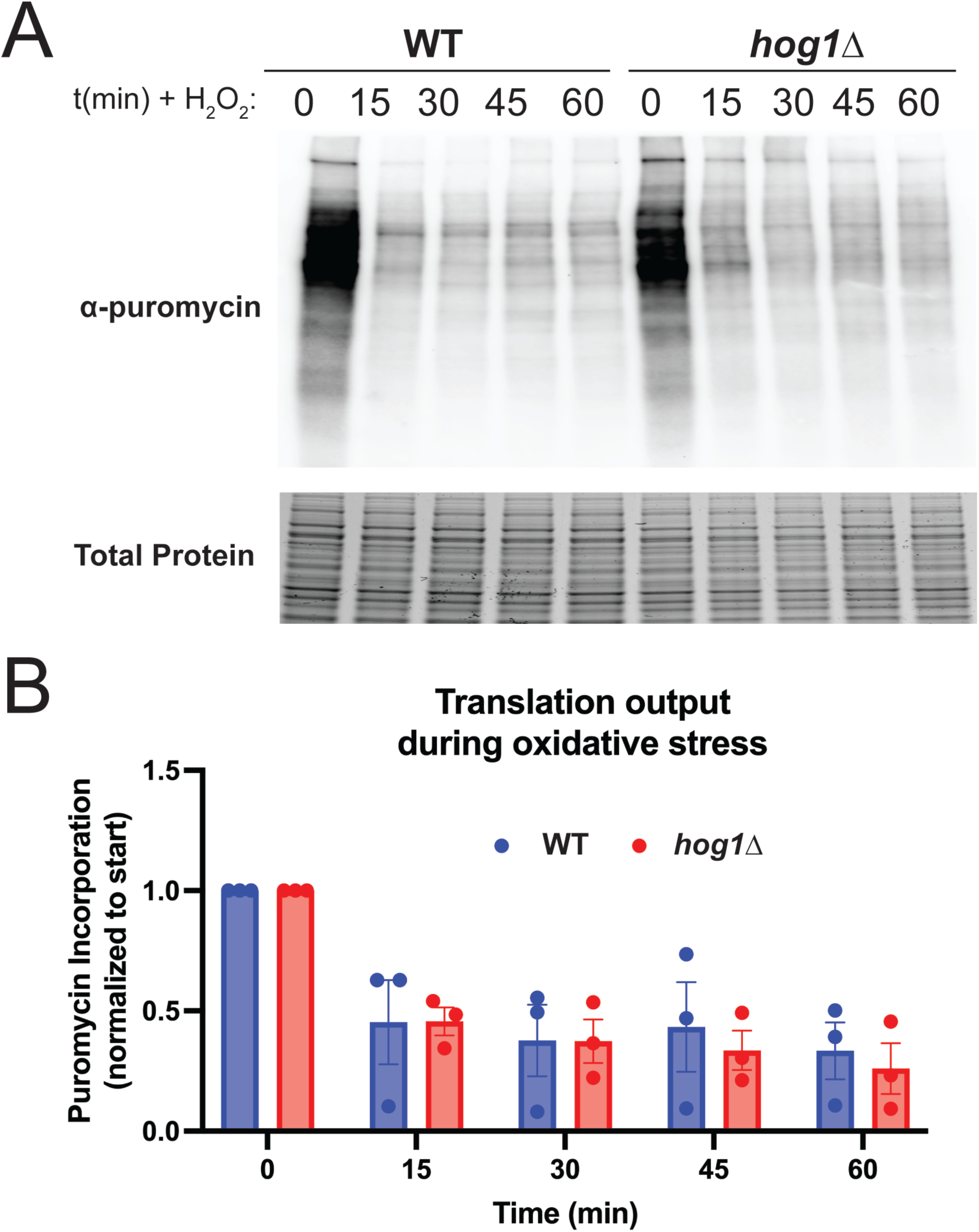
Puromycin incorporation assay in WT and *hog1*Δ during oxidative stress. (A-B) WT and *hog1*Δ were grown to midlogarithmic phase and subjected to 2mM H_2_O_2_ for the indicated amounts of time. Puromycin was added and incorporated for 10 minutes prior to harvest of each time point. (A) Protein lysates were prepared and probed for puromycin using an anti-puromycin antibody. Total protein is shown as a loading control. (B) Results from three biological replicates were quantified by densitometry, using total protein signal as the denominator. Then signal for each strain was normalized to the untreated (*t* = 0) sample.

**Table S1:**
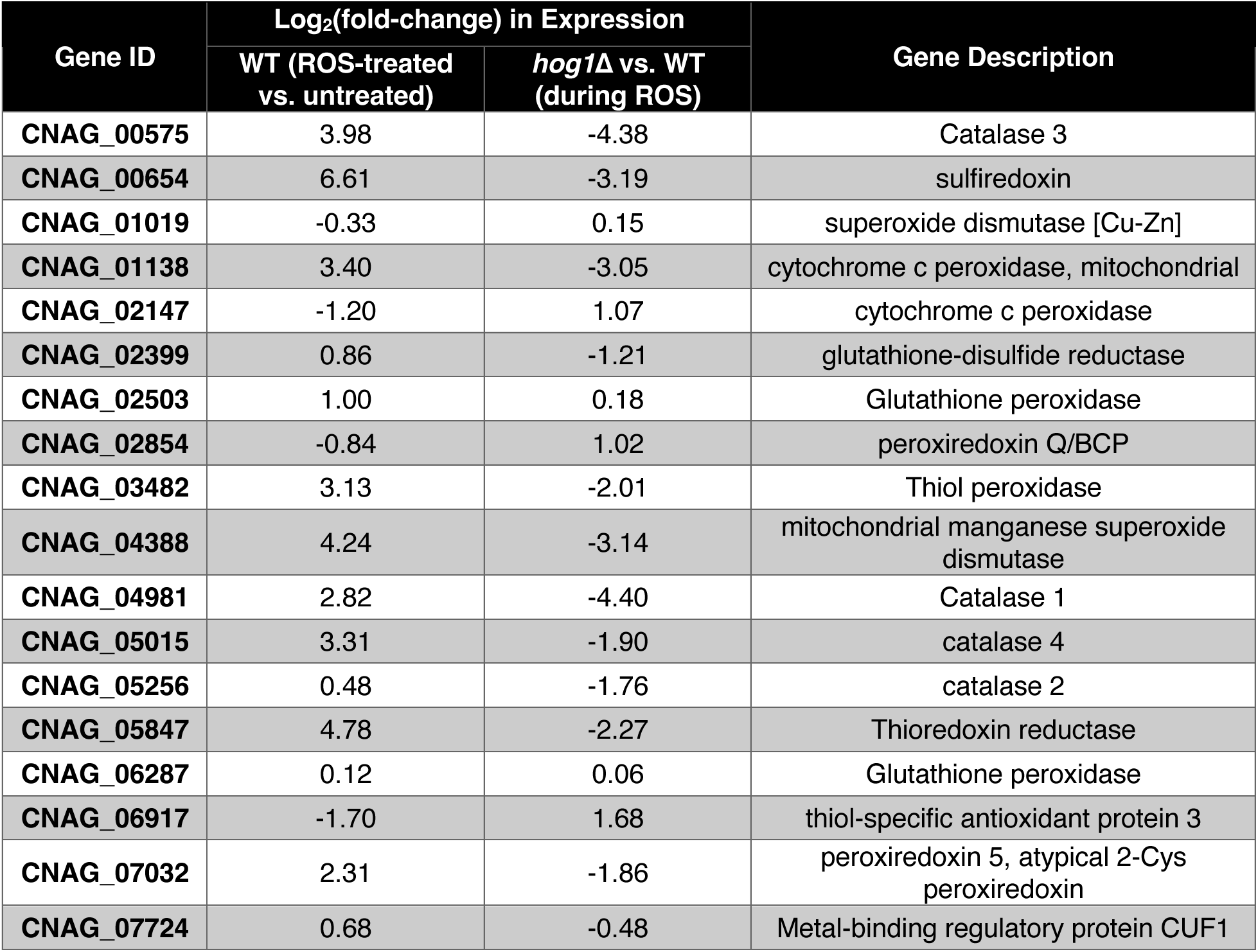
Expression of antioxidants in the WT oxidative stress response and differential expression in *hog1*Δ. Transcripts listed under the GO term “antioxidant activity” are shown. Log2(fold-change) in expression was calculated using DEseq2 for the indicated pairwise comparison of samples.

**Table S2:**
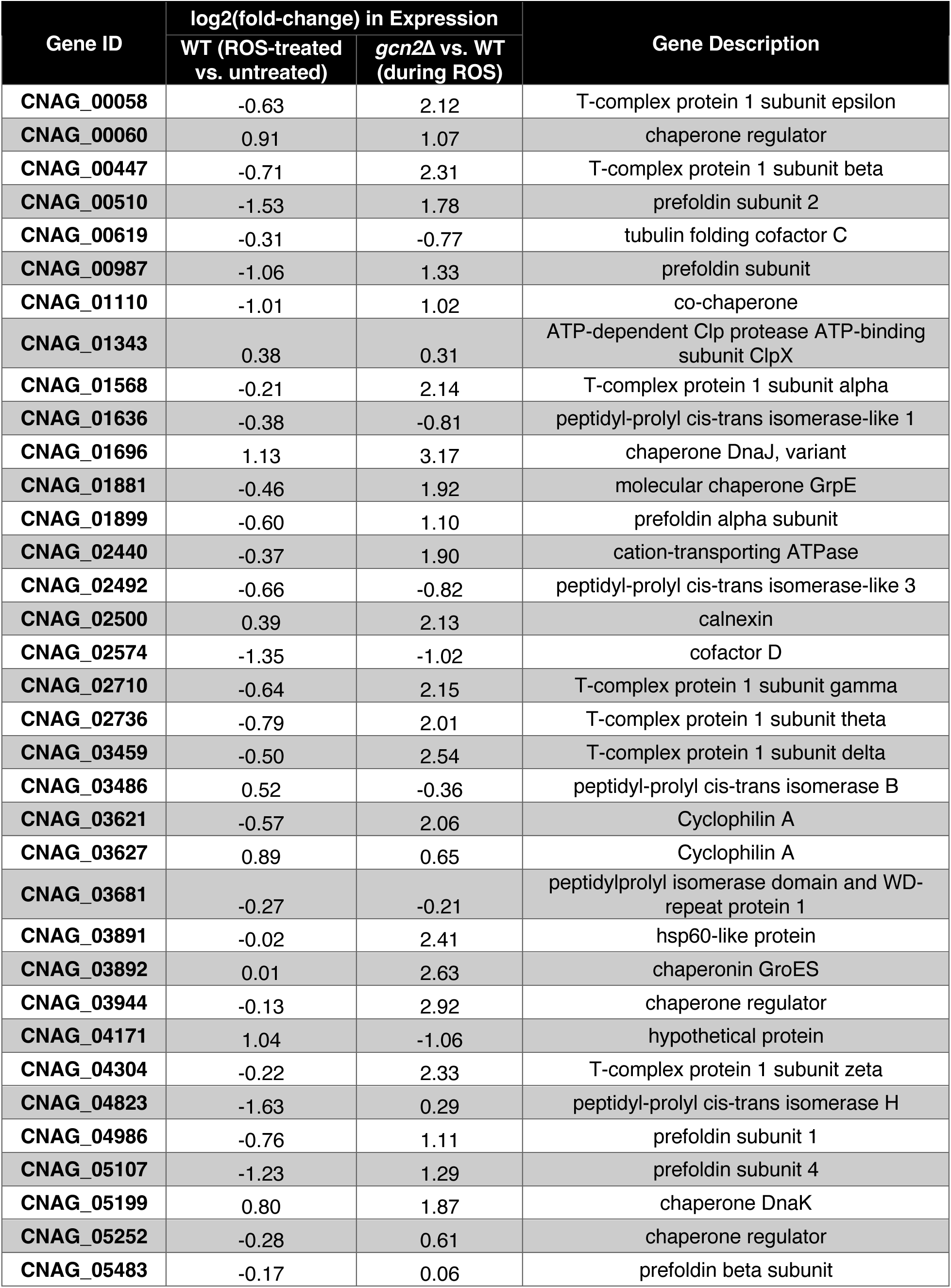
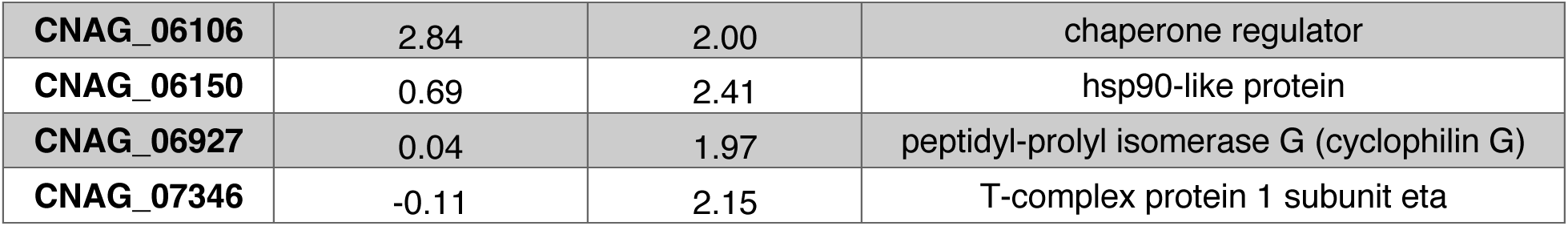
Expression of protein folding genes in the WT oxidative stress response and differential expression in *gcn2*Δ. Transcripts listed under the GO term “protein folding” are shown. Log2(fold-change) in expression was calculated using DESeq2 for the indicated pairwise comparison of samples.

**Table S3:**
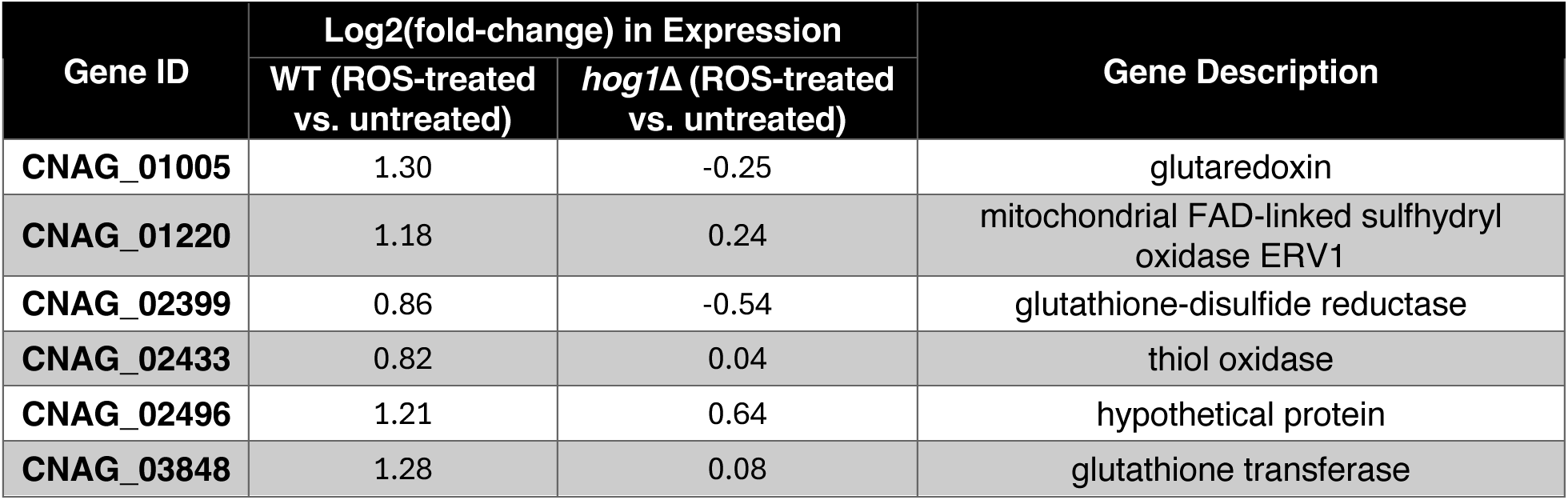
Expression of protein disulfide reductases that failed to be upregulated in *hog1*Δ ROS response. Transcripts that were upregulated in the WT ROS response, but not the *hog1*Δ ROS response, and fall under the GO term “protein-disulfide reductase” are shown. Log2(fold-change) in expression was calculated using DEseq2 for the response of the indicated strains.

**Table S4:**
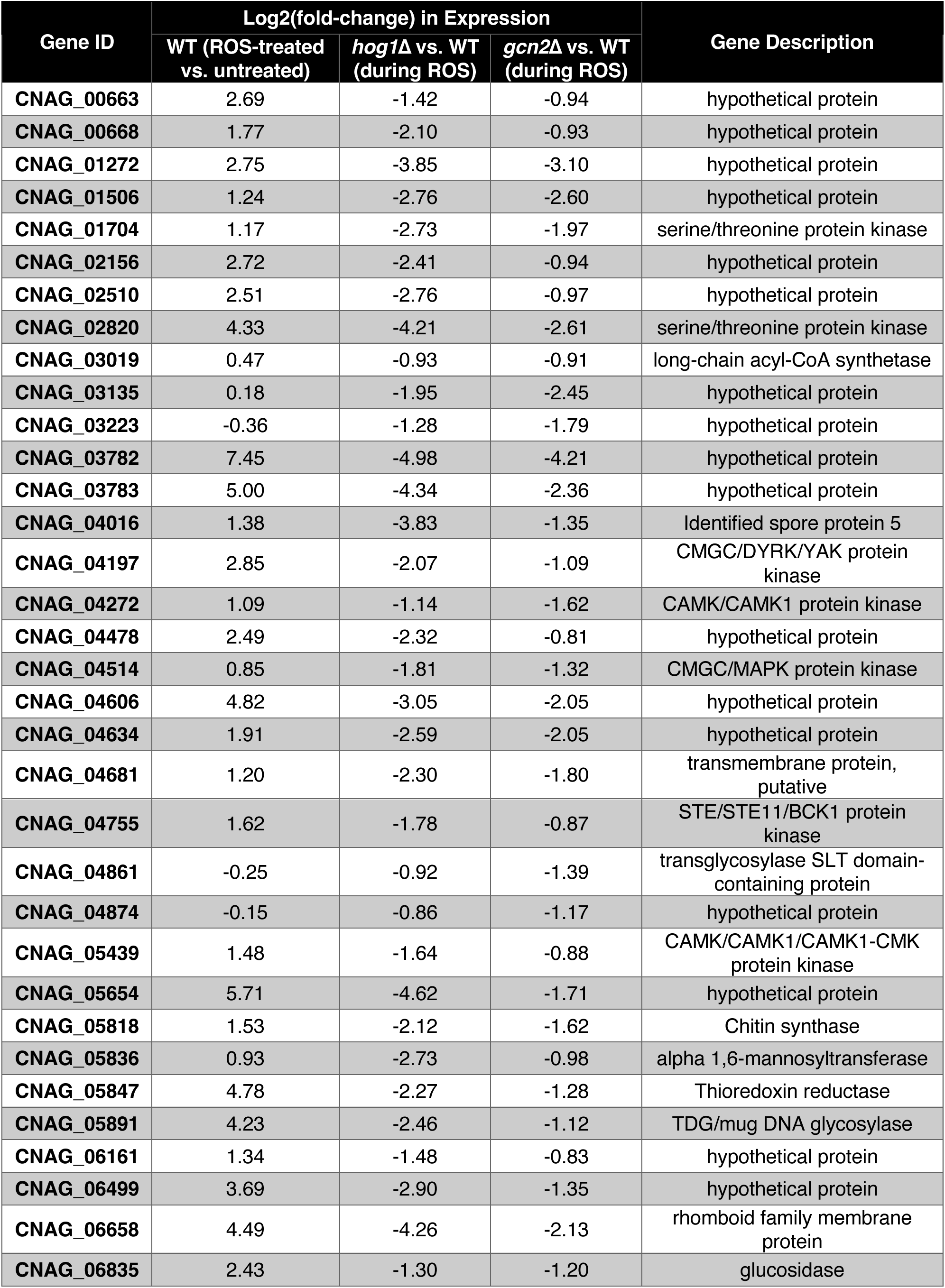
Stress response factors with lower expression in *hog1*Δ and *gcn2*Δ, relative to WT, during oxidative stress. Transcripts with decreased expression in *hog1*Δ and *gcn2*Δ fall under the GO term “cellular response to stress” are shown. Log2(fold-change) in expression was calculated using DEseq2 for the indicated pairwise comparisons of samples.

**Table S5:** Selected targets with decreased expression in *hog1*Δ and increased expression in *gcn2*Δ, relative to WT during oxidative stress. Selected transcripts from the list of 226 transcripts with decreased expression in *hog1*Δ and increased expression in *gcn2*Δ, relative to WT during oxidative stress. We selected transcripts with putative function in redox processes and detoxification. Log2(fold-change) was calculated using DEseq2 for the indicated pairwise comparisons of samples.

| Gene ID | Log2(fold-change) in Expression |  |  | Gene Description |
| --- | --- | --- | --- | --- |
|  | WT (ROS-treated vs. untreated) | *hog1Δ* vs. WT (during ROS) | *gcn2Δ* vs. WT (during ROS) |  |
| CNAG_00121 | 2.02 | -1.79 | 0.90 | glycerol-3-phosphate dehydrogenase (NAD()) |
| CNAG_00154 | 2.83 | -2.47 | 1.11 | oxidoreductase |
| CNAG_01584 | 4.00 | -4.49 | 1.76 | hydrolase |
| CNAG_01604 | 3.41 | -1.33 | 2.28 | TIGR01458 family HAD hydrolase |
| CNAG_01605 | 3.57 | -3.26 | 1.18 | rossman fold oxidoreductase |
| CNAG_02519 | 2.70 | -0.97 | 1.53 | mitochondria fission 1 protein |
| CNAG_05573 | 1.50 | -1.17 | 0.86 | cytochrome c oxidase assembly protein subunit 17 |
| CNAG_06868 | 5.04 | -4.22 | 1.39 | Phosphopyruvate hydratase |
| CNAG_06880 | 1.99 | -1.40 | 1.19 | mitochondrial protein |

**Table S6:** Selected targets with uniquely lower expression in *hog1*Δ*gcn2*Δ, with respect to WT during oxidative stress. Selected transcripts from the list of 314 transcripts with decreased expression in *hog1*Δ*gcn2*Δ, with respect to WT during oxidative stress, that did not reach thresholds for differential expression in single knockouts. We selected transcripts involved in metabolic processes, ion transport, and regulation of the cytoskeleton, which appeared to be most prevalent in this gene list. The log2(fold-change) for the indicated pairwise comparisons is shown.

| Gene ID | Log2(fold-change) in Expression |  |  |  |  | Gene Description |
| --- | --- | --- | --- | --- | --- | --- |
|  | WT (ROS-treated vs. untreated) | (ROS-treated mutant vs. ROS-treated WT) |  |  |  |  |
|  |  | *hog1*Δ | *gcn2*Δ | *hog1*Δ*gcn2*Δ #9 | *hog1*Δ*gcn2*Δ #11 |  |
| CNAG_00098 | 2.13 | -0.55 | -0.47 | -1.39 | -1.31 | palmitoyl-protein thioesterase |
| CNAG_00318 | 0.32 | -0.49 | 0.12 | -0.98 | -0.87 | arp2/3 complex 16 kda subunit |
| CNAG_00450 | 2.28 | 0.71 | -0.76 | -1.64 | -1.47 | 3-isopropylmalate dehydrogenase |
| CNAG_00749 | 3.03 | 0.08 | -1.38 | -4.60 | -4.95 | alternative sulfate transporter |
| CNAG_00905 | 1.10 | -0.69 | 0.08 | -0.90 | -1.31 | MFS transporter |
| CNAG_01489 | 0.28 | -0.54 | -0.16 | -1.41 | -1.28 | maltose O-acetyltransferase |
| CNAG_03245 | 1.69 | -0.75 | -0.03 | -2.03 | -1.94 | glucose-6-phosphate dehydrogenase |
| CNAG_03666 | 0.87 | -0.35 | -0.64 | -0.88 | -1.22 | acyl-CoA dehydrogenase |
| CNAG_03920 | 1.98 | 0.41 | -0.60 | -1.51 | -1.43 | isocitrate dehydrogenase, NADP-dependent |
| CNAG_04013 | -0.40 | -0.70 | 0.55 | -0.97 | -1.10 | potassium channel protein |
| CNAG_04765 | 1.64 | -0.59 | 0.23 | -1.11 | -0.94 | ARP2/3 complex 21 kDa subunit |
| CNAG_04898 | 0.97 | -0.35 | -0.15 | -0.95 | -0.94 | MFS transporter |
| CNAG_05737 | 2.24 | -0.63 | -0.53 | -1.86 | -1.77 | oligopeptide transporter 8 |
| CNAG_06890 | 0.91 | -0.18 | -0.26 | -1.33 | -1.79 | membrane transporter |
| CNAG_07582 | 1.11 | -0.10 | 0.18 | -1.13 | -1.27 | dynein light chain roadblock-type |

**Table S7:** Translation initiation factors with dysregulated expression in *hog1*Δ, relative to WT during ROS. Among transcripts upregulated in *hog1*Δ vs. WT during oxidative stress, transcripts with names containing “translation initiation factor” were selected. Log2(fold-change) in expression was calculated using DESeq2.

| Gene ID | Log2(fold-change) expression<br>in *hog1Δ* vs. WT (during<br>oxidative stress) | Gene Description |
| --- | --- | --- |
| CNAG_00580 | 0.90 | translation initiation factor IF-2 |
| CNAG_01428 | 1.49 | Translation initiation factor 5A |
| CNAG_01660 | 0.92 | translation initiation factor eif-2b subunit<br>alpha |
| CNAG_01709 | -1.54 | translation initiation factor 5 |
| CNAG_02128 | 1.19 | translation initiation factor 3 subunit J |
| CNAG_02657 | 1.04 | translation initiation factor 3 subunit G |
| CNAG_04022 | -0.86 | translation initiation factor |
| CNAG_04260 | 1.45 | translation initiation factor eIF-2B subunit<br>gamma |
| CNAG_05007 | 1.11 | translation initiation factor eIF-2B subunit<br>beta |
| CNAG_05843 | -1.25 | translation initiation factor 4E |
| CNAG_07628 | 0.96 | translation initiation factor 3 subunit C |
| CNAG_07778 | 0.92 | translation initiation factor 2 subunit 1 |

## Notes

### Competing Interest Statement

The authors have declared no competing interest.

